# Diminished Stimulus-Evoked Activity During Sustained Attention Without Behavioural Cost in ADHD

**DOI:** 10.64898/2026.07.09.737359

**Authors:** Elaine Pinggal, David Chapman, Amy Q. Huynh, Alessia Ruyant-Belabbas, Redmond G. O’Connell, Jennifer Windt, Sean P.A. Drummond, Tim J. Silk, Mark A. Bellgrove, Thomas Andrillon

**Affiliations:** School of Psychological Sciences, Monash University, Clayton, VIC, 3800, Australia; Turner Institute for Brain and Mental Health & School of Psychological Sciences, Faculty of Medicine, Nursing, and Health Sciences, Monash University, Melbourne, VIC, 3800, Australia; Monash Alfred Psychiatry Research Centre (MAPrc), Melbourne, VIC, 3004, Australia; Paris Brain Institute, Sorbonne Université, Institut National de la Santé et de la Recherche Médicale (Inserm)-Centre National de la Recherche Scientifique (CNRS), Paris 75013, France; Trinity College Institute of Neuroscience and School of Psychology, Trinity College Dublin, Dublin D02 PN40, Ireland; School of Philosophical, Historical, and International Studies, Monash University, Melbourne, VIC, 3800, Australia; Monash Centre for Consciousness and Contemplative Studies, Monash University, Melbourne, VIC, 3800, Australia; Deakin Institute for Lifespan Health & Development and the School of Psychology, Deakin University, Burwood, 3125, Australia; Developmental Imaging, Murdoch Children’s Research Institute, Melbourne, VIC, 3052, Australia

**Author notes:** **Corresponding author:** Elaine Pinggal,; Thomas Andrillon. These authors contributed equally (Mark A. Bellgrove, Thomas Andrillon).

**Keywords:** Electroencephalogram, attention-deficit/hyperactivity disorder, slow waves, local sleep, sustained attention, arousal

## Abstract

Attention-Deficit/Hyperactivity Disorder (ADHD) is often accompanied by attentional challenges, sleep difficulties and increased reports of daytime sleepiness, indicating potential links between attention and arousal mechanisms.

Electroencephalographic (EEG) studies have shown that sleep-like wake slow wave (SW) activity during wakefulness, characterised by reduced cortical activity, corresponds with periods of inattention in both ADHD and neurotypical populations. However, it remains unclear whether group differences in waking SWs are consistent across different types of sustained attention tasks or may be context-dependent. The present study compared sustained attention performance and wake SW activity between adults with and without ADHD during a continuous visual target detection task.

EEG data were collected while adults with (*n* = 52) and without ADHD (*n* = 49) completed a sustained attention paradigm. The task presented a continuous stream of rotating (anti-clockwise or clockwise) black-and-white checkerboard stimuli, and participants were required to detect infrequent targets that were marginally longer in duration than non-targets. Behavioural and neural measures (power spectra, steady-state visually evoked potentials (SSVEP), event-related potentials and SW density during wake) were analysed.

No significant group differences were observed for task performance and wake SW activity. However, electrophysiological analyses revealed the ADHD group showed reduced P300, elevated beta-band power, and lower SSVEPs to target stimuli compared with the neurotypical group.

These neural differences in the context of comparable performance suggest compensatory cortical recruitment in ADHD. The absence of SW differences further supports this, suggesting comparable performance and maintained arousal may reflect successful neural compensation in ADHD in this cognitive task.

**Significance Statement:** Individuals with ADHD commonly experience attention differences, but emerging evidence suggests these difficulties may depend on task context. Using clinical diagnostic interviews, medication washout, and Bayesian statistics, we found strong evidence for comparable performance between ADHD and neurotypical adults on a sustained attention task. Notably, slow wave activity during wakefulness did not differ between groups, consistent with maintained arousal. Nevertheless, differences in brain activity, including altered target processing, increased cortical activation and reduced visual entrainment, suggested compensatory neural recruitment in ADHD. This context-dependent pattern has important theoretical and clinical implications: identifying task features that challenge or support performance in ADHD can reveal how individuals compensate neurally, and guide interventions that leverage task features to support attention in ADHD.

## Introduction

Despite decades of research, the precise neural mechanisms underlying attentional difficulties in Attention Deficit/Hyperactivity Disorder (ADHD) remain elusive. ADHD affects approximately 2.5% of adults and is characterised by pervasive symptoms of inattention, hyperactivity, and/or impulsivity that significantly impact cognitive performance and everyday functioning (Faraone et al., 2024; May et al., 2023). While behavioural studies consistently demonstrate that individuals with ADHD exhibit greater response variability and higher error rates during sustained attention tasks (Bellgrove et al., 2005; Bozhilova et al., 2020; Helfer et al., 2020; Madiouni et al., 2020; Pinggal et al., 2026), understanding why these fluctuations occur has proven challenging.

ADHD symptoms vary with cognitive demands (Fortenbaugh et al., 2017; Raffaelli et al., 2025), yet research typically employs similar protocols, such as variations of the continuous performance task (CPT) or the Sustained Attention to Response Task (SART) (Robertson et al., 1997). Task demands can modulate arousal and the emergence of sleep-like neural activity during wakefulness, a relationship that may partly account for attentional variability across paradigms. These mechanisms, however, remain poorly understood, limiting our understanding of these processes in ADHD.

Sleep and attention regulation provide compelling insights into neural bases of ADHD. Sustained attention enables individuals to maintain focus during everyday activities, fluctuating with sleep quality (Gobin et al., 2015), motivation (Esterman et al., 2016), and task engagement (Smallwood et al., 2004). Sleep deprivation in neurotypical individuals produces behavioural symptoms resembling ADHD’s core characteristics: impulsivity, distractibility, and sustained attention difficulties (Beebe, 2011; Choong et al., 2025; Drummond et al., 2006, 2012; Gruber et al., 2012; Krause et al., 2017; Saletin et al., 2019), alongside disrupted prefrontal network activity, also observed in ADHD (Cortese et al., 2012; Czisch et al., 2012; Krause et al., 2017; Ma et al., 2015; Saletin et al., 2019). These parallels suggest sleep processes during wakefulness may critically influence ADHD symptom expression.

Sleep disturbances affect 25-55% of individuals with ADHD (Cortese et al., 2013; Hvolby, 2015), including nocturnal awakening, restless sleep, parasomnias, and delayed or shortened sleep (Bioulac et al., 2015; Bjorvatn et al., 2017; Hvolby, 2015). They also report increased daytime sleepiness and exhibit elevated frontal delta activity during rest (Helfer et al., 2020). These findings suggest that sleep-like neural activity, which naturally intrudes during wakefulness even in neurotypical individuals (Andrillon et al., 2021; Pinggal et al., 2022), may appear with unique characteristics or increased frequency in ADHD, potentially influencing cognitive function.

Local sleep, wherein sleep-like SWs appear in specific brain regions during wakefulness (Andrillon et al., 2019; Vyazovskiy et al., 2011), has been associated with fluctuations in sustained attention across populations. These high-amplitude SWs represent brief reductions in neuronal activity and increase with extended wakefulness or task engagement (Andrillon et al., 2021; Bernardi et al., 2015; Hung et al., 2013; Steriade, 2005; Vyazovskiy et al., 2011). Their regional occurrence predicts distinct cognitive outcomes: frontal SW correlate with impulsivity (false alarms or faster responses), whereas parietal SW are associated with sluggishness (missed trials or slower responses) (Andrillon et al., 2021; Pinggal et al., 2022). These differential neural patterns suggest local sleep intrusions act as a functional switch, temporarily disrupting specific brain networks and corresponding cognitive functions. While this relationship has been established in neurotypical individuals, our recent work provided evidence that increased SW may also contribute to attentional difficulties in ADHD (Pinggal et al., 2026), though the generalisation of this result remains to be tested across different paradigms.

The present study examined whether sleep-like SWs in wakefulness were associated with sustained attention decrements in individuals with and without ADHD. The Continuous Temporal Expectation Task (CTET), a sustained attention task previously untested in an ADHD population, differs from conventional detection-based paradigms in its reliance on temporal expectation rather than simple stimulus detection, providing a novel context in which to examine SW-related attentional lapses. With error rates typically around 30–40% in neurotypical adults (Connell et al., 2009), the CTET represents a moderately demanding paradigm whose suitability for detecting group differences in ADHD is yet to be established. We hypothesised that adults with ADHD would exhibit: (1) lower task performance; (2) elevated sleep-like SW compared with neurotypical adults; and (3) a negative correlation between SWs and performance.

## Materials and Methods

### Participants

One hundred and one adult participants (*N* = 101; 52 with ADHD and 49 neurotypical individuals) were recruited in accordance with Monash University ethical guidelines (project numbers 17839 and 29393). Neurotypical adults had no history of neuropsychiatric disorders and scored below the clinical thresholds for ADHD on both the Adult ADHD Self-Report Scale for DSM-5 (ASRS-5; threshold:14) (Ustun et al., 2017) and the ADHD Index subscale from the Conners’ Adult ADHD Rating Scales (CAARS; threshold: 65) (Conners et al., 1999). One neurotypical participant did not complete the CAARS but scored below the ASRS-5 threshold and reported no history of ADHD. Two participants, one from each group, were excluded from our analyses (incomplete dataset: 1; participation withdrawal: 1). The final sample consisted of 99 participants (N = 51 ADHD and N = 48 neurotypical adults, see Table 1 for demographics and clinical characteristics).

**Table 1.** Demographics and Questionnaire Data.

| | ADHD ($N = 51$) | Neurotypical ($N = 48$) | *p*-Value |
| --- | --- | --- | --- |
| Age (years, mean (SD)) | 29.7 (10.4) | 25.1 (7.4) | .01 |
| Sex (% females) | 66.7% | 64.6% | .95 |
| Education level (years, mean (SD)) | 16.3 (3.3) | 15.5 (2.9) | .23 |
| Other lifetime psychiatric diagnoses |  |  |  |
| Depression | 58.8% | 2.0% |  |
| Any Anxiety Disorder | 60.8% | 0% |  |
| Post Traumatic Stress Disorder | 3.9% | 0% |  |
| Personality Disorder | 2.0% | 0% |  |
| Bipolar Disorder | 3.9% | 0% |  |
| Autism | 11.8% | 0% |  |
| Eating Disorder | 11.8% | 0% |  |
| Obsessive Compulsive Disorder | 5.9% | 0% |  |
| Substance Dependence Disorder | 2.0% | 2.0% |  |
| Dyslexia | 3.9% | 0% |  |
| Tourette's Syndrome | 2.0% | 0% |  |
| Questionnaire data |  |  |  |
| **Adult ADHD Self-Report Scale (ASRS-5) (mean (SD))** | 16.7 (3.5) | 6.6 (2.6) | < .001 |
| **Conners' Adult ADHD Rating Scale (CAARS)** |  |  |  |
| **DSM-IV Inattentive Symptoms** (Subscale E, mean (SD)) | 81.8 (8.5) | 50.7 (11.6) | < .001 |
| **DSM-IV Hyperactive-Impulsive Symptoms** (Subscale F, mean (SD)) | 69.7 (9.9) | 43.9 (8.0) | < .001 |
| **DSM-IV Total ADHD Symptoms** (Subscale G, mean (SD)) | 80.3 (8.9) | 47.2 (10.8) | < .001 |
| **ADHD Index** (Subscale H, mean (SD)) | 71.0 (9.0) | 47.5 (9.5) | < .001 |
| **ADHD presentation (DIVA diagnosis)** |  |  |  |
| Combined | 74.5% |  |  |
| Inattentive | 25.5% |  |  |
| Hyperactive-impulsive | 0% |  |  |

**Note.** ADHD = Attention Deficit/Hyperactivity Disorder. Higher scores on the ASRS-5 (total score) and CAARS subscales indicate greater ADHD-related symptom severity.

Clinical interviews (Diagnostic Interview for ADHD in Adults (DIVA 2.0)) were conducted by a trained psychiatrist to confirm ADHD diagnoses and determine the patients’ ADHD presentation: “inattentive” (N=13), “hyperactive/impulsive” (N=0) or “combined” (N=38). Participants completed online questionnaires assessing demographics, ADHD symptoms (CAARS, ASRS-5, Wender Utah Rating Scale), depression and anxiety (subscales from the Inventory of Depression and Anxiety Symptoms (IDAS-II)), and substance use (Alcohol Use Disorders Identification Test, Drug Use Disorders Identification Test). In addition, participants completed the National Adult Reading Test (NART) in person to estimate verbal IQ. The results from the DIVA, CAARS and ASRS-5 questionnaires are presented in Table 1, while findings from the remaining measures are provided in Supplementary Table 1.

Except for the twelve participants (N = 5 Inattentive and N = 7 Combined) who were not taking medication at the time of recruitment and testing, all participants with ADHD interrupted their treatment 72 hours before testing (details of participants’ ADHD medication are provided in Supplementary Table 2).

### Study Design

Participants completed an online battery of questionnaires assessing ADHD symptoms, mood and substance use. Those in the ADHD group completed a structured clinical interview (DIVA 2.0) and observed a three day ADHD medication washout period prior to the experimental session (see *Participants* section and Supplementary Table 2).

All participants attended a session at Monash University’s Clayton campus. They first completed the NART to estimate verbal IQ, followed by eight blocks of the Continuous Temporal Expectancy Task (CTET) while their brain activity was recorded using electroencephalography (EEG). Participants were seated in a Faraday room and fitted with a 64-channel EEG cap. A two minute resting state was recorded before and after the task, though these data are not included in the present analysis.

### Continuous Temporal Expectancy Task (CTET)

The CTET assesses sustained attention through a visual monitoring paradigm (O’Connell et al., 2009). Participants viewed a centrally presented patterned stimuli against a grey background featuring a square divided into a 10 by 10 grid of diagonally split black and white squares (Figure 1a). Each stimulus frame randomly rotated 90° clockwise or counter-clockwise, creating four distinct arrangements. A central white cross was used to direct the gaze of participants to minimise eye movements and each pattern flickered at 25 Hz to generate steady-state visually evoked potentials (SSVEP). Participants were instructed to monitor, detect and respond to target stimuli by pressing a button. The stimuli were presented at two durations: 800 ms (standard, non-target) or 1120 ms (target), with 7–15 non-target stimuli pseudo-randomly inserted between targets, resulting in 5.6–12 s intervals between each target presentation (see Supplementary Figure 1 for an animated depiction of the task).

**Figure 1.**
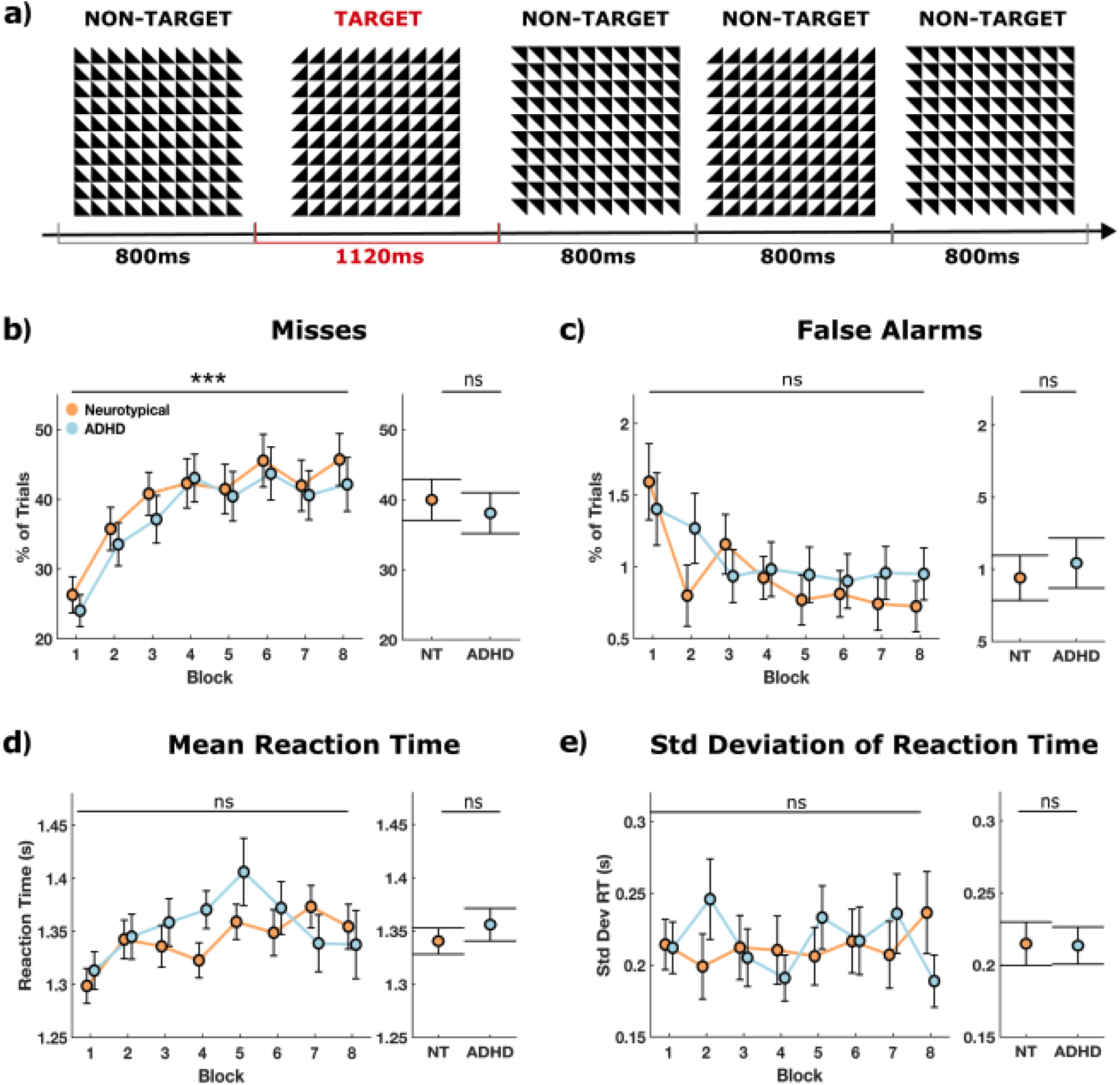
Experimental Protocol and Continuous Temporal Expectancy Task (CTET) Performance ***Note*. a.** Participants completed eight blocks of the CTET, a sustained attention task in which they monitored a continuous stream of black-and-white checkerboard stimuli that rotated anti-clockwise or clockwise. They were instructed to press a button in response to target stimuli (1120 ms), which were identical in appearance to non-targets (800 ms) but differed in presentation duration. **b. – e.** Performance metrics including: **b.** misses, **c**. false alarms, **d**. mean reaction time, and **e.** standard deviation of reaction time. The left panels show the mean performance of each group (Neurotypicals in orange and ADHD in blue) across blocks (N = 8 blocks) with error bars representing the standard error. The right panels show the overall mean and standard error by group. Stars indicate the strength of the block effect (left panels) and group differences (right panels) as determined by Bayesian linear mixed-effect models (ns: BF₁₀ <⅓, evidence for null; ***BF₁₀ > 30, strong evidence).

Prior to the main experiment, participants completed a practice session consisting of two blocks: the first with three targets among 25 standard stimuli, and the second incorporating the 25 Hz flicker. Participants were required to achieve 100% accuracy before completing the main experiment. The main experiment consisted of eight experimental blocks, each containing 225 frames, 18–22 target presentations, and lasting approximately three minutes.

### Behavioural Data

Behavioural performance was analysed by computing the proportion of missed targets and false alarms. A target stimulus was considered missed if no response was registered between 800 ms after target onset and 1600 ms after target offset (reaction time range: 0.8–2.72 s). This range was chosen as target stimuli are longer in duration (1120 vs. 800 ms), allowing participants to correctly detect a stimulus when it is presented for longer than 800 ms. We then waited for the equivalent of two standard stimuli (1600 ms) after the end of the target stimulus to ensure that participants did not perform a late response. False alarms were defined as responses to non-target stimuli, specifically when no target stimulus had occurred in the two preceding trials (no target stimuli within the last 1600 ms). This approach prevented any overlap between the definition of misses and false alarms. Reaction times for target trials were calculated from stimulus onset.

Performance indices for each CTET block were extracted, including the percentage of missed targets, false alarms, average reaction times for correctly detected targets (in seconds) and reaction time variability (standard deviation of reaction times).

### EEG

#### Data Acquisition and Preprocessing

Participants sat in a dim room where high-density EEG data were recorded using an actiCAP with 64 active scalp electrodes with a BrainAmp DC system (Brain Products, Munich, Germany), referenced to FCz, and sampled at 500 Hz. Participants completed a two-minute resting state before and after the CTET during which they were instructed to fixate on a white cross in the centre of the screen while blinking naturally as needed.

Preprocessing of EEG data was performed using custom MATLAB scripts with the FieldTrip toolbox (Oostenveld et al., 2011). The continuous EEG data were either segmented following the task blocks (8 blocks per recording section, −1 s to +1 s before block onset and after block offset) or into stimulus-locked epochs (−0.5 s to +1.8 s relative to stimulus onset).

The trial-based data segments were preprocessed as follows: baseline correction (−0.2 to 0 s), bandpass filtering using a fourth-order Butterworth filter (0.1–40 Hz), notch filtering (50, 100 and 150 Hz, Discrete Fourier Transform filter) to remove power line noise, and re-referencing all EEG channels to the average of all electrodes. Baseline correction was applied within a pre-stimulus interval. Following preprocessing, the data were down-sampled to 250 Hz.

Prior to further analysis, data quality was assessed through visual inspection to identify and remove noisy channels and trials containing artefacts that were initially flagged for high deviance from the rest of the signal. Flagged channels were interpolated using a weighted neighbour approach to reconstruct the missing data. Independent Component Analysis (ICA) was then performed to identify and remove components associated with ocular and cardiac artefacts and were removed based on visual inspection. To extract the ICA components, we changed the filtering of the EEG from 0.1–40 Hz to 1–40 Hz.

In addition to trial-based analysis, data were also segmented into longer epochs defined by the onset and offset of each block (−1s before and +1s after block onset and offset). The same preprocessing pipeline was applied to these block-level data, including identical filtering parameters, re-referencing and down-sampling procedures. Additionally, the same noisy channels and ICA components identified during the trial-level preprocessing were removed from the block-level data to ensure consistency across analyses.

We used block-epoched data for behaviour, power spectrum and slow wave analysis, and trial-epoched data for ERP analysis.

#### Event Related Potentials (ERPs)

Trial-epoched EEG data were re-referenced to the average of TP9 and TP10 electrodes (located near the mastoid regions) and baseline corrected (−0.2 to 0 s). Due to the task design – where infrequent target stimuli were longer in duration (1120 ms) than frequent non-targets (800 ms) – we focused on offset-locked ERPs to obtain neural responses following the end of stimulus presentation. Trials were aligned to the time point immediately following stimulus offset, and ERPs were computed separately for target and non-target trials.

#### Spectral analysis

The power spectral density (PSD) was computed for each electrode using the block-epoched data. Spectral analysis was performed using Welch’s method with a 60-second window and 50% overlap between successive windows, independent of the preprocessing epoch length. The frequency resolution was set to 0.1 Hz across the 1–40 Hz range. Prior to spectral analysis, each block was baseline-corrected by subtracting the mean amplitude across the time series for each electrode. Power spectra were computed independently for each block and electrode, then averaged across blocks to obtain subject-level estimates. The resulting power values were log-transformed to normalise the distribution. Additionally, signal-to-noise ratio (SNR) was calculated for each frequency bin using a minimum distance parameter of 0.5 Hz, with specific extraction of the 25 Hz SNR value (corresponding to the SSVEP frequency) for further analysis.

#### Sleep-like slow waves

Sleep-like SW were identified using established algorithms to detect SW in sleep (Riedner et al., 2007) and wakefulness (Andrillon et al., 2021; Pinggal et al., 2022, 2026). EEG signals were re-referenced to TP9 and TP10 (located near the mastoid regions) and the mean removed from each channel. All SW were first identified by detecting consecutive negative zero-crossings (start and end of individual waves respectively). For each wave, key parameters were extracted including amplitude (peak-to-peak; PTP), duration (time difference between the start and end), downward and upward slopes, and frequency (1/duration). Waves were discarded if they exhibited frequencies above 7 Hz, PTP amplitudes exceeding 150 µV, or a positive peak above 75 µV.

To focus on large amplitude SW, which have been shown to share properties with sleep SW, we selected individual waves with the highest PTP amplitude. However, given the between-subject design, we used a common threshold across the entire sample at the electrode level. To do so, we applied the same method as a previous study comparing ADHD and neurotypical adults during a SART (Pinggal et al., 2026). First, electrode-specific amplitude thresholds were established from the neurotypical group. We derived slow wave thresholds by calculating the 80th percentile of PTP amplitude values for each electrode in each neurotypical participant across all detected waves. These values were then averaged across the neurotypical group for each electrode to obtain a single electrode-wise threshold. Second, for all participants (both ADHD and neurotypical), SW exceeding this neurotypical-derived threshold were classified as high-amplitude “sleep-like” SW. Consistent with previous studies (Andrillon et al., 2021; Bernardi et al., 2015; Hung et al., 2013; Pinggal et al., 2022, 2026; Quercia et al., 2018; Vyazovskiy et al., 2011), we restricted the detection of SW within the δ–θ frequency range ([1–7] Hz).

### Statistical analyses

#### Demographics

To compare demographics between the ADHD and neurotypical groups, we conducted *t*-tests (age, education, ASRS-5, CAARS) and a chi-square test (sex; *p-*values in Table 1).

#### Behaviour

Analysis of performance on the CTET was conducted using a Bayesian linear mixed-effects (LME) modelling approach in *R* (brm function from the brms package). As such, *p*-values are not reported in this section. Instead, we present posterior estimates, 95% credible intervals, and Bayes Factors (BF) to quantify the evidence for or against additional predictors to the model. The BF indicates whether the observed correlation is more likely under the alternative hypothesis (i.e., a correlation exists) versus the null hypothesis (i.e., no correlation exists). The alternative hypothesis assumes correlations follow a Cauchy distribution where 50% of the distribution falls between −0.33 and 0.33. In this framework, a *BF₁_0_* greater than 3 provides evidence supporting the alternative hypothesis, whereas values less than ⅓ favour the null hypothesis that no meaningful correlation exists.

#### Event Related Potentials (ERPs)

To examine group differences in offset-locked ERPs at electrode Pz, we computed a difference wave for each participant by subtracting ERPs to non-target trials from ERPs to target trials. We then conducted a timepoint-by-timepoint one-way ANOVA with group (ADHD vs neurotypical) as the between-subjects factor to identify time windows where the amplitude of the ERP difference wave significantly differed between groups.

To determine whether participants showed a difference in the P300 component, we computed a mean ERP difference score at electrode Pz by subtracting non-target from target trials and averaging across a 0.15–0.45 s post-stimulus offset window, which encompasses the expected P300 peak. For each group, one-sample *t*-tests were used to assess whether the mean difference wave was significantly different from zero. To evaluate P300 group differences, an independent sample *t*-test was performed comparing mean difference amplitudes between the ADHD and neurotypical groups.

To identify electrode clusters showing significant group differences in ERP difference wave amplitude, a cluster-based permutation test was applied across all electrodes and time-points (Maris & Oostenveld, 2007; α = .05, two-tailed Monte-Carlo α = .025). Group differences were visualised using a topographic map at the peak timepoint of the largest significant cluster, defined as the timepoint with the maximum summed t-statistic across cluster electrodes.

#### Spectral Analysis

Group differences in PSD across all electrodes and frequencies (1–40 Hz) were assessed using a cluster-based permutation test (Maris & Oostenveld, 2007). For the SSVEP analysis, electrode Oz was selected based on its highest SNR for the 25 Hz frequency-tagged response, consistent with the expected occipital distribution of visual steady-state responses. Group differences in SSVEP amplitude at 25 Hz were assessed using an independent sample *t*-test at Oz. Group differences in 25 Hz SNR across all electrodes were further assessed using a cluster-based permutation test (Maris & Oostenveld, 2007; 5000 permutations, cluster α = .05, two-tailed Monte-Carlo α = .025) and visualised using a topographic map.

#### Slow Waves

To examine the effects of block, group and age on SW density, we conducted Bayesian model comparisons using LME models. We compared nested models with different combinations of fixed effects (block, group and age) while including participant ID as a random effect to account for repeated measures. Model comparisons were evaluated using BF to quantify the evidence for including additional predictors, with the same criteria described above for the behavioural analyses.

To further determine whether there was a difference in SW density across the scalp between groups, we implemented one LME per electrode with SW density as the dependent variable, group, block and age as fixed effects, and participant ID as a random effect. For each electrode, we extracted *t*-statistics and *p*-values for the group effect (ADHD vs neurotypical). To identify electrode clusters showing significant group differences in SW density, we applied a cluster-based permutation test across all electrodes (Maris & Oostenveld, 2007; 5000 permutations, cluster α = .05, two-tailed Monte-Carlo α = .025).

## Code Accessibility

All code used for this study are available at: https://github.com/andrillon/CTET_ADHD

## Results

### Demographics

Demographic characteristics were compared between the ADHD and neurotypical groups. Independent sample *t*-tests indicated that participants in the ADHD group were significantly older than those in the neurotypical group (*t*(90.30) = −2.56, *p* = .012); therefore, we evaluated whether including age improved model fit in the LME analyses and included it as a covariate in models where it significantly improved model fit.

There was no significant difference between groups in years of education (*t*(94.18) = −1.21, *p* = .229). A chi-square test of independence showed that distribution of sex did not significantly differ between groups (χ² = 2.05, *p* = .951).

### Behaviour

A series of Bayesian model comparisons were conducted to evaluate the effects of block, group and age on CTET performance measures: misses, false alarms, reaction times and reaction time variability (see Table 2). For misses, the best fitting model included block number as a fixed effect and subject identity as a random intercept. In contrast, for false alarms, reaction time and reaction time variability, the best fitting model included only subject identity as a random intercept. All best fitting models excluded group and age as predictors, with Bayes Factors providing strong evidence against the inclusion of these predictors (see Table 2). This suggests there were no meaningful group and age differences in CTET performance.

**Table 2.** Bayesian model comparisons for the inclusion of block, group and age across CTET performance measures.

| Performance | Predictor | $BF_{01}$ | $BF_{10}$ |
| --- | --- | --- | --- |
| **Misses** |  |  |  |
|  | **Block** | **$3.21 \times 10^{-21}$** | **$2.92 \times 10^{20}$** |
|  | Group | 22.29 | 0.04 |
| | Age | 453.62 | $2.28 \times 10^{-3}$ |
| **False alarms** |  |  |  |
| | Block | $3.00 \times 10^8$ | $8.85 \times 10^{-9}$ |
| | Group | 389.80 | $2.55 \times 10^{-3}$ |
| | Age | 7434.57 | $1.30 \times 10^{-4}$ |
| **Reaction Time** |  |  |  |
| | Block | $1.12 \times 10^9$ | $9.09 \times 10^{-10}$ |
| | Group | 38.63 | $2.62 \times 10^{-2}$ |
| | Age | 791.90 | $1.25 \times 10^{-3}$ |
| **Reaction Time Variability** |  |  |  |
| | Block | $4.20 \times 10^{12}$ | $2.34 \times 10^{-13}$ |
| | Group | 51.50 | $1.98 \times 10^{-2}$ |
| | Age | 950.37 | $1.07 \times 10^{-3}$ |

**Note.** **Bold** = predictors that were added to the model. $BF_{01}$ represents the Bayes factor in favour of the null (reduced) model over the alternative (full) model, quantifying evidence against including a predictor. Conversely, $BF_{10}$ is the inverse ( $1/BF_{01}$ ) and quantifies the evidence for including the predictor. Values of $BF_{01} > 3$ (or $BF_{10} < 1/3$ ) indicate moderate-to-strong evidence for the null; values of $BF_{10} > 3$ (or $BF_{01} < 1/3$ ) indicate moderate-to-strong support for the alternative. Values between $1/3$ and 3 are considered inconclusive.

Results revealed strong evidence that miss rates were modulated by block (*BF₁_0_ =* 2.92 × 10^20^) and increased over time as shown in Figure 1b. For false alarms, reaction time and reaction time variability, model comparisons provided strong evidence against the inclusion of block number and group effects, with Bayes factors strongly favouring the null model (see Table 2). To assess whether incorporating individual symptom severity improved model fit, additional models including the CAARS ADHD Index and total ADHD symptom scores were compared against the winning null model; results are presented in Supplementary Table 3. In all of these models, the evidence at hand supported or strongly supported the exclusion of these variables as predictors of performance. Additionally, exploratory analysis using a 65% performance threshold on block one, excluding participants who missed more than 35% of targets in the first block, is presented in Supplementary Figure 2. These additional analyses also found evidence for an absence of group effects across performance metrics.

### ERP

The task stimuli evoked marked ERPs with a difference between NoGo and Go stimuli corresponding to a classical P300 at stimulus offset. To examine group differences in offset-locked ERP responses, we first completed a timepoint-by-timepoint one-way ANOVA with group (ADHD vs neurotypical) as the between-subjects factor on the ERP difference wave (target minus non-target) at electrode Pz. No time windows showed a significant difference in ERP amplitude between groups (Figure 2a).

**Figure 2.**
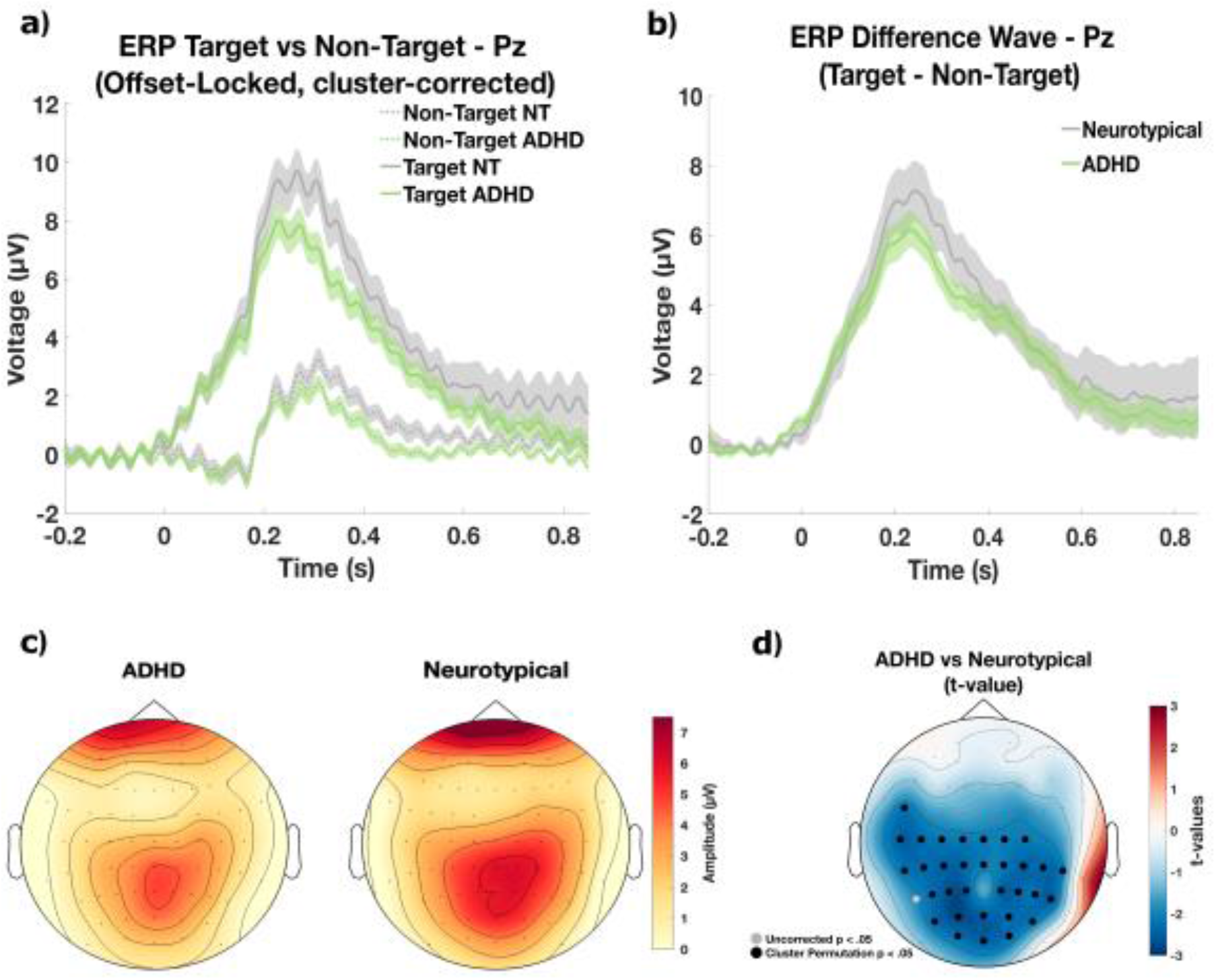
Event Related Potential: Group Comparisons ***Note*. a.** Event related potential (ERP) waves offset-locked at electrode Pz comparing target (continuous lines) and non-targets (dashed lines) for neurotypical (NT; grey) and ADHD (green) groups. Shaded areas around each line represent standard error of the mean (SEM). No significant group differences were observed. **b.** ERP difference wave (target minus non-target) for each group. The shaded areas around each line represent SEM. **c.** Topographical maps of the ERP difference wave amplitude averaged across 0.15–0.45 s post-stimulus offset for the ADHD (left) and NT (right) groups. **d.** Topography of group differences (ADHD minus NT) in ERP difference wave amplitude at the cluster peak timepoint (0.261 s). *T*-values from a cluster-based permutation test are shown. Black dots indicate electrodes belonging to the significant negative cluster at the peak timepoint (overall cluster: 0.129–0.345 s, *p* = .040, cluster-corrected), reflecting lower P300 amplitude in the ADHD group, while grey dots indicate significant electrodes with uncorrected *p* < .05.

To evaluate P300 (or P3b) responses more directly, we computed mean amplitudes within the 0.15–0.45 s post-stimulus offset window at electrode Pz (Figure 2b), i.e. the standard time and electrode at which the P300 is typically observed. The target–non-target difference wave was significantly greater than zero in both the ADHD (*t*(50) = 11.57, *p* < .001) and neurotypical group (*t*(47) = 7.20, *p* < .001). However, there was no significant difference in P300 amplitude between groups (*t*(97) = 0.99, *p =* .33).

To further examine the spatial distribution of group differences across all electrodes, a cluster-based permutation test was applied across the full ERP window and across electrodes. A significant negative cluster was found, ranging 0.129–0.345 s post-stimulus offset (peak: 0.261 s, *p* = .040), which overlapped substantially with the a priori P300 window (0.15–0.45 s) but did not include Pz. This indicates that the ADHD group showed broadly lower ERP difference wave amplitude compared to the neurotypical group across this window (Figure 2d). No significant positive clusters were observed.

### Power Spectrum

Power spectral analysis compared PSD (1–40 Hz) across all electrodes between groups, with Fz shown for illustrative purposes (Figure 3a). A cluster-based permutation test revealed a significant positive cluster ranging 13.3–21.3 Hz (peak 18.3 Hz, *p* = .035), indicating elevated beta-band power in the ADHD group relative to the neurotypical group across frontal, central and parietal electrodes (Figure 3b). No significant negative clusters were observed.

**Figure 3.**
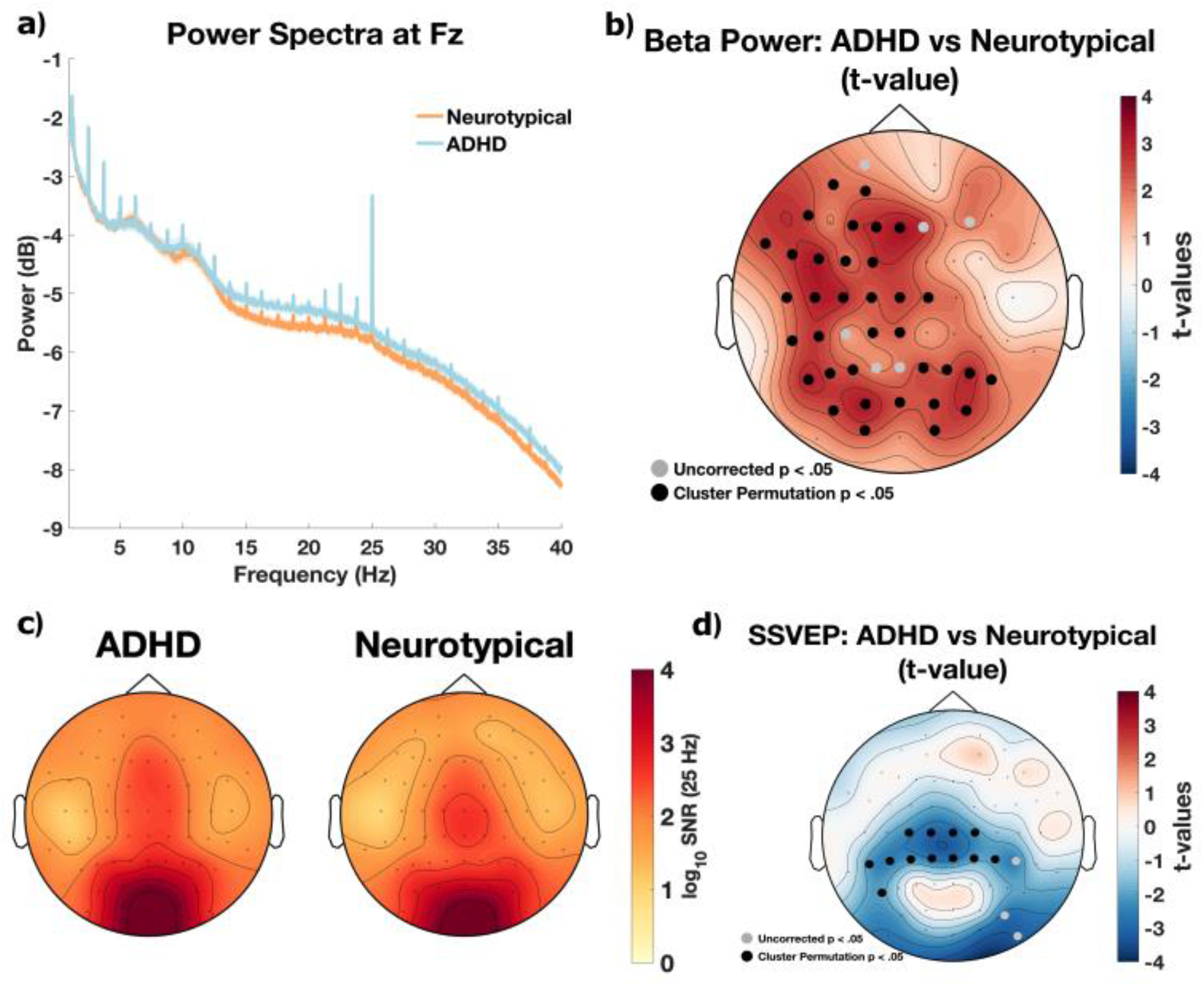
Power Spectra and Steady-State Visually Evoked Potentials Response Differences Between Groups ***Note.* a.** Power spectral density at electrode Fz across 1–40 Hz comparing neurotypical (NT; orange) and ADHD (blue). The shaded areas around each line represent SEM. **b.** Topography of group differences in beta-band power (ADHD minus NT) at the cluster peak frequency (18.3 Hz). Black dots indicate cluster-corrected significant electrodes, reflecting higher beta power in the ADHD group, while grey dots indicate significant electrodes with uncorrected *p* < .05. **c.** Topographical maps of the signal to noise ratio (SNR) across the scalp for the ADHD (left) and NT (right) groups. **d.** SSVEP group difference topography (ADHD minus NT) at 25 Hz, reflecting lower SSVEP amplitude in the ADHD group over central and parietal electrodes. Black dots indicate cluster-corrected significant electrodes (*p* = .016); grey dots indicate electrodes with uncorrected *p* < .05.

The SSVEP analysis revealed a clear frequency tag at 25 Hz in both groups. Topographical analysis showed both groups exhibited typical high activity in occipital regions (Figure 3c), consistent with the entrainment of activity of early visual cortices with SSVEP paradigms. An independent-sample *t*-test at electrode Oz revealed no significant group difference in 25 Hz SNR (*t*(97) = −1.56, *p* = .123, Cohen’s d = −0.31). However, a cluster-based permutation test across all electrodes identified a significant negative cluster (*p* = .016), indicating lower SSVEP amplitude in the ADHD group over central and parietal electrodes (Figure 3d).

### Slow Wave Density During Wakefulness

We completed a series of Bayesian model comparisons to evaluate the effects of block, group and age on SW density (model comparison results shown in Table 3). Results revealed strong evidence that slow wave density was modulated by block (BF_01_ = 4.81 × 10^-16^; *BF₁_0_ =* 2.50 × 10^15^). As shown in Figure 4c, SW density differed across blocks, with ADHD participants tending towards higher values than neurotypical participants and a slight downward shift over time. When comparing models that included block alone to one including both block and group, the estimated Bayes Factors (BF_01_ = 1.16; *BF₁_0_* = 0.82) indicated inconclusive evidence in favour of either model, suggesting that adding group did not substantially improve model fit. In contrast, when comparing a model including block alone to one including both block and age, the Bayes Factor (BF_01_ = 3.72 × 10^-5^; *BF₁_0_* = 24812.47) provided very strong evidence in favour of including age, indicating that age substantially improved model fit for predicting SW density. The estimate for age (β = −0.51, 95% CI = [-0.71, −0.32]) suggests that SW density decreased with increasing age.

**Figure 4.**
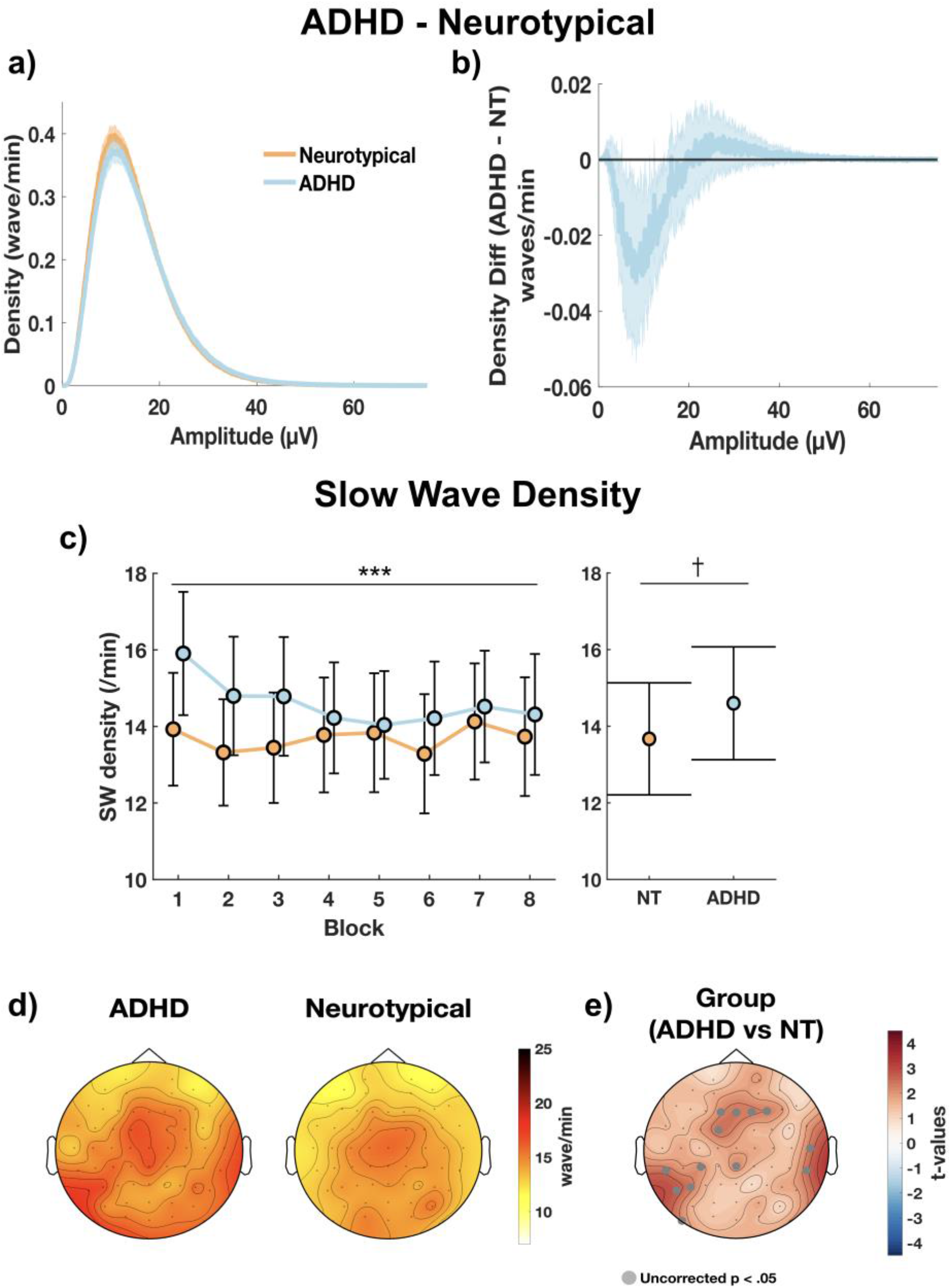
Slow Wave Density Across Groups ***Note.* a.** Distribution of slow wave amplitude (waves per minute) showing that neurotypical participants (NT; orange) had a higher density of low-amplitude slow waves, while ADHD participants (blue) showed a higher density of high-amplitude slow waves. **b.** Difference wave (ADHD minus NT) with shaded standard error of the mean (SEM) showing the shift in amplitude toward higher-amplitude slow waves in ADHD. **c.** The left panel depicts the mean slow wave density of each group (NT in orange, ADHD in blue) across blocks, with error bars representing the standard error. The right panel shows the overall mean slow wave density and standard error by group. Symbols indicate the strength of the block effect on slow wave density (left panel) and group differences (right panel), as determined by Bayesian linear mixed-effect models (†: ⅓ ≤ BF₁₀ < 3, inconclusive; ***: BF₁₀ > 30, strong evidence). **d.** Topographical maps of the slow wave density distribution across the scalp for the ADHD and NT groups. **e.** Statistical comparison showing *t*-values for the group effect (ADHD vs NT) extracted from electrode-wise linear mixed-effect models. Grey dots indicate electrodes with uncorrected *p* < .05. No electrodes survived cluster-based permutation correction for multiple comparisons.

**Table 3.**
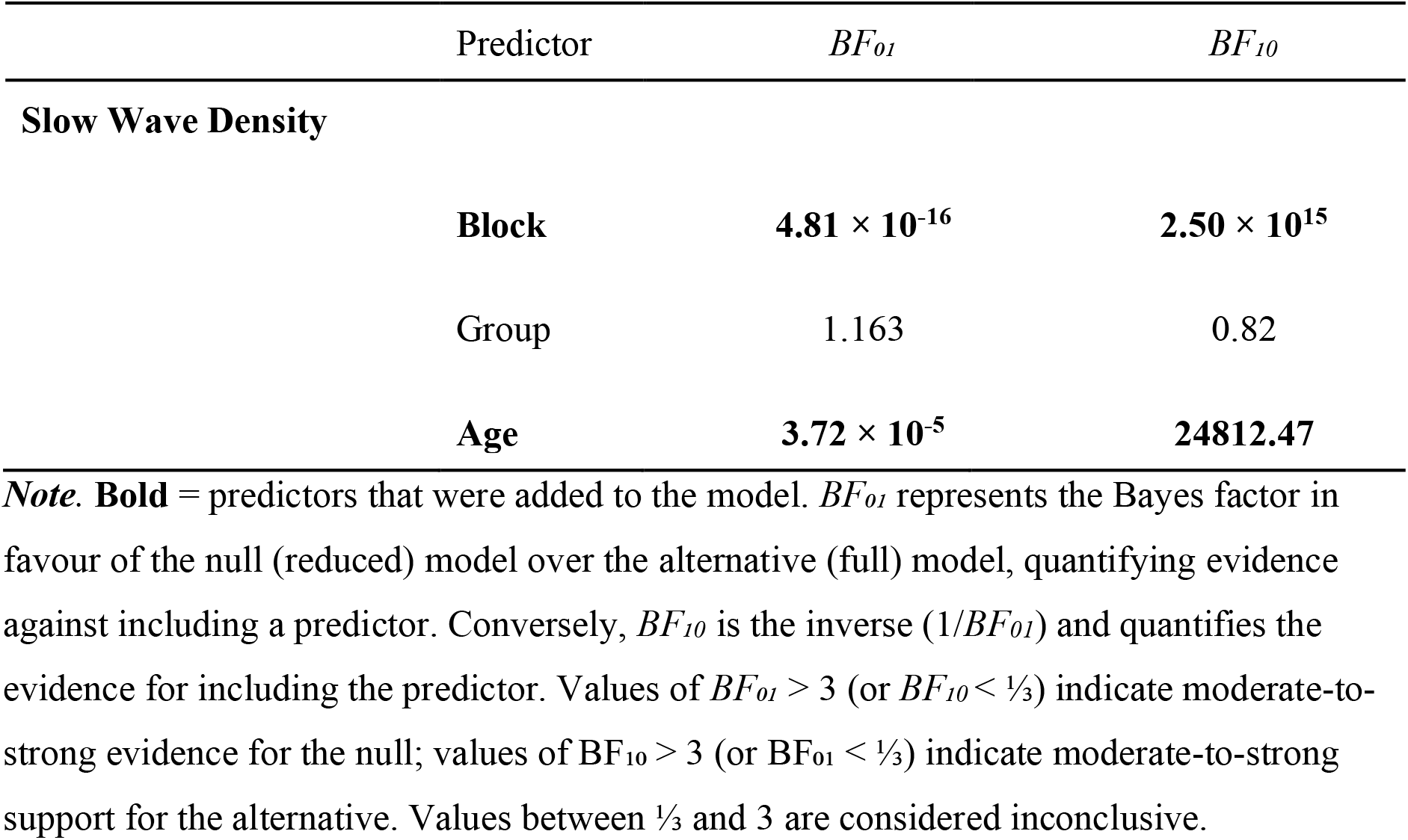
Bayesian model comparisons for the inclusion of block, group and age on slow wave density.

| Predictor | $BF_{01}$ | $BF_{10}$ |
| --- | --- | --- |
| **Slow Wave Density** |  |  |
| **Block** | **$4.81 \times 10^{-16}$** | **$2.50 \times 10^{15}$** |
| Group | 1.163 | 0.82 |
| **Age** | **$3.72 \times 10^{-5}$** | **24812.47** |

*Note.* **Bold** = predictors that were added to the model. $BF_{01}$ represents the Bayes factor in favour of the null (reduced) model over the alternative (full) model, quantifying evidence against including a predictor. Conversely, $BF_{10}$ is the inverse ( $1/BF_{01}$ ) and quantifies the evidence for including the predictor. Values of $BF_{01} > 3$ (or $BF_{10} < 1/3$ ) indicate moderate-to-strong evidence for the null; values of $BF_{10} > 3$ (or $BF_{01} < 1/3$ ) indicate moderate-to-strong support for the alternative. Values between $1/3$ and 3 are considered inconclusive.

To investigate whether SW density differed across the scalp between groups, we fitted separate LME models for each electrode, with SW density as the dependent variable, and group, block and age as fixed effects. Participant ID was included as a random intercept to account for repeated measures. Uncorrected group effects appeared most prominently over frontal, centro-parietal and left posterior electrodes. However, no electrodes survived cluster-based permutation correction for multiple comparisons (Figure 4e).

## Discussion

### Comparable CTET Performance: Potential Explanations

Contrary to expectations, the ADHD and neurotypical groups performed comparably on the CTET. Bayes Factors strongly supported this null finding despite consistent evidence reported previously of greater response variability and higher error rates in ADHD during sustained attention tasks (Bellgrove et al., 2005; Bozhilova et al., 2020; Helfer et al., 2020; Madiouni et al., 2020; Pinggal et al., 2026). These results may reflect task features that preserve attentional performance in ADHD.

The CTET’s temporal structure may have inadvertently facilitated performance in the ADHD group. The continuous and regular rotation of standard stimuli in the task provided a rhythmic background that participants could potentially entrain to automatically, without requiring deliberate sustained attention (Coull & Nobre, 2008). Indeed, some participants reported noticing a rhythm to the rotating squares, indicating awareness and potential use of these temporal cues. Critically, while adults with ADHD show greater variability in self-generated motor rhythms, their ability to synchronise movements to external auditory rhythmic cues remains comparable to neurotypical individuals (Kliger Amrani & Zion Golumbic, 2020). In the present study, individuals with ADHD may have similarly entrained their attention to this rhythmic background, leveraging their preserved ability to synchronise to external rhythms to partly compensate for sustained attention difficulties. It should be noted, however, that the high overall miss rates across both groups (∼40%) suggests the task was challenging for participants, which limits the extent to which this fully accounts for the comparable performance found. Nonetheless, these findings suggest rhythmic structure and automatic entrainment as potential avenues for supporting attention in ADHD.

Beyond rhythmic entrainment, the CTET may have engaged different cognitive processes than those typically assessed in, for example, SART-based paradigms. Unlike the SART, which requires responses to frequent Go trials while withholding responses to rare NoGo trials, the CTET requires responses only to infrequent targets. This reversal creates greater cognitive load and sustained attention demands in the SART, particularly for response inhibition, a domain typically more challenging for individuals with ADHD. This likely explains the more pronounced group differences and greater time-on task effects observed with the SART (Bozhilova et al., 2020; Machida et al., 2022; Pinggal et al., 2022, 2026; Robertson et al., 1997; Van Den Driessche et al., 2017).

Moreover, the slight decrease in global SW density, rather than the expected increase, suggests participants did not experience the progressive build-up of sleep pressure and mental fatigue that characterises more cognitively demanding sustained attention paradigms (Dijk, 2009; Lazar et al., 2015; Lo et al., 2012; Riedner et al., 2007). The decrease in false alarms across early task blocks may also reflect ongoing task familiarisation, suggesting that learning during experimental blocks may have obscured expected vigilance decrements.

Additionally, during the EEG set up, many participants in the ADHD group expressed enthusiasm for contributing to ADHD research, suggesting heightened intrinsic motivation that may have contributed to comparable performance beyond the monetary incentive provided to neurotypical participants (Esterman et al., 2016). This motivational factor, combined with the task design features described above, may have collectively contributed to comparable performance between groups.

### Electrophysiological Findings

Both groups showed the expected P300 component (250–400 ms) (Polich, 2007). A cluster-based permutation test revealed a significant negative cluster ranging 0.129–0.345 s post-stimulus offset, indicating that the ADHD group showed broadly reduced P300 amplitude compared to the neurotypical group across posterior and central electrodes (Figure 2d). However, this difference was not reflected at electrode Pz alone, suggesting the effect may be spatially distributed rather than localised to a single canonical electrode. These results align with the literature where individuals with ADHD typically exhibit significantly reduced P300 amplitude during target detection and decision-making tasks compared to neurotypical individuals (Biabani et al., 2025; Bozhilova et al., 2020; Kaiser et al., 2020; Szuromi et al., 2011), which has been attributed to greater cognitive effort and engagement of additional neural resources to achieve comparable performance (Szuromi et al., 2011).

Power spectral analysis (1–40 Hz) revealed elevated beta power in the ADHD group relative to the neurotypical group across frontal, central and parietal electrodes. This mirrors trends reported by Loo et al. (2009), who similarly observed elevated beta power alongside comparable task performance in adults with ADHD, and interpreted this as reflecting compensatory cortical recruitment. The present findings extend this, providing more robust evidence that increased cortical activation in ADHD may serve to achieve comparable performance. While resting-state spectral power may not reliably differentiate adults with ADHD from neurotypical individuals (Liechti et al., 2013; Markovska-Simoska & Pop-Jordanova, 2017), the present findings suggest that task-based paradigms may be more sensitive to spectral differences in adult ADHD.

Both groups showed a clear entrainment to the standard stimulus presentation at 25 Hz, with the SSVEP frequency tag maximal over occipital electrodes, as expected. However, the ADHD group showed significantly lower SSVEP responses over central and parietal electrodes, while no significant difference was observed at electrode Oz specifically. Since SSVEP amplitude can reflect sustained visual attention to flickering stimuli (Morgan et al., 1996; Müller et al., 1998), the lower SSVEP amplitude in the ADHD group over central and parietal electrodes suggests differences in attentional engagement rather than solely early visual processing.

Finally, we examined an EEG marker of sleepiness to investigate potential arousal differences between groups, in line with our previous work on sleep intrusions during wakefulness and attentional lapses in individuals with or without ADHD (Andrillon et al., 2021; Pinggal et al., 2026). SW density decreased across blocks, with strong Bayesian evidence for this time-on-task effect. This is surprising given that SW density typically increases over time during sustained attention tasks, reflecting increasing mental fatigue and reduced alertness (Andrillon et al., 2019; Hung et al., 2013; Pinggal et al., 2026; Vyazovskiy et al., 2011). Model comparisons provided inconclusive evidence for group differences (*BF₁_0_* = 0.82), suggesting group membership did not substantially improve model fit. However, age significantly improved model fit, with older individuals showing lower SW density; a pattern consistent with age-related decreases during both sleep (Crowley, 2002) and wakefulness (Vlahou et al., 2014). Since the ADHD group was significantly older, this age-related decrease may have obscured potential group differences, as any elevation in the ADHD group could have been offset by age-related reductions. Uncorrected topographical maps suggested higher SW density in the ADHD group over frontal, centro-parietal, and left posterior electrodes compared to the neurotypical group, though no electrodes survived cluster-based permutation correction. These uncorrected patterns thus remain exploratory and do not provide evidence for reliable group differences in SW density within the current paradigm.

Notably, these results contrast with the higher SW density observed in adults with ADHD during a SART (Pinggal et al., 2026). The absence of a significant group difference in SW density in the CTET alongside intact task performance may be related to the nature of the task and an increase in engagement, motivation, and/or arousal in the ADHD group. The *decrease* in SW during the CTET, contrasting with the typical *increase* during a SART, supports this interpretation. Taken together, the absence of group differences in both behaviour and SW density supports the notion that attentional differences in ADHD may be context-dependent, with task features shaping not only performance but also underlying arousal mechanisms. Future studies investigating group differences in wake SW patterns should perhaps employ more cognitively demanding paradigms or examine how task structure (e.g., temporal predictability, cognitive demands) modulates SW density, while ensuring better age-matching between groups.

### Future Directions

Our findings suggest that rhythmic or predictable task structures may provide cognitive support for individuals with ADHD, as evidenced by comparable performance between groups. Future research could systematically manipulate temporal predictability and rhythmic elements to identify the conditions under which they support attention, and whether individuals with ADHD derive greater benefit from temporal cues compared to neurotypical individuals. Rather than avoiding predictable task structure in ADHD research, researchers could leverage rhythmic or predictable elements to study compensatory entrainment strategies more directly, or explore the utility of their incorporation into cognitive training programs for ADHD. Measuring or manipulating participants’ motivation could also provide key insights on individual and group differences.

## Conclusion

This study examined sustained attention and SW density between adults with ADHD and neurotypical adults completing the CTET. Contrary to expectations, no reliable behavioural or SW group differences were found, though several neural differences emerged. Individuals with ADHD showed reduced P300 amplitude, elevated beta power, and lower SSVEP amplitude compared to neurotypical adults, suggesting compensatory cortical recruitment to maintain comparable performance. The temporal structure and rhythmic elements of the task may have also facilitated this compensation, supporting attention in the ADHD group. Together, these findings highlight the context-dependent nature of ADHD-related attentional differences and suggest that temporal predictability may represent a promising avenue for cognitive support interventions in ADHD.

## Supplementary Tables

**Supplementary Table 1.**
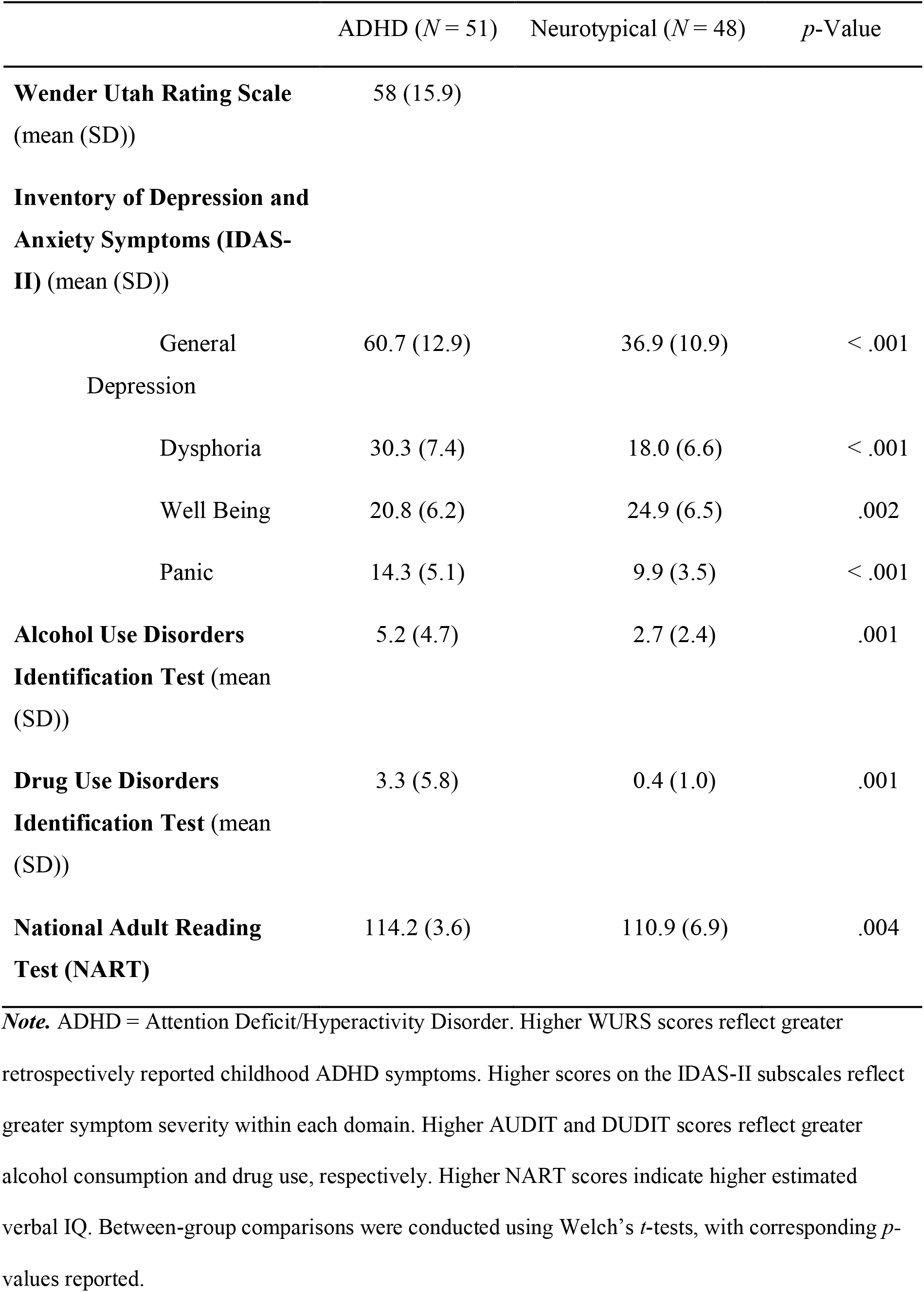
Demographics and Questionnaire Data.

**Supplementary Table 2.**
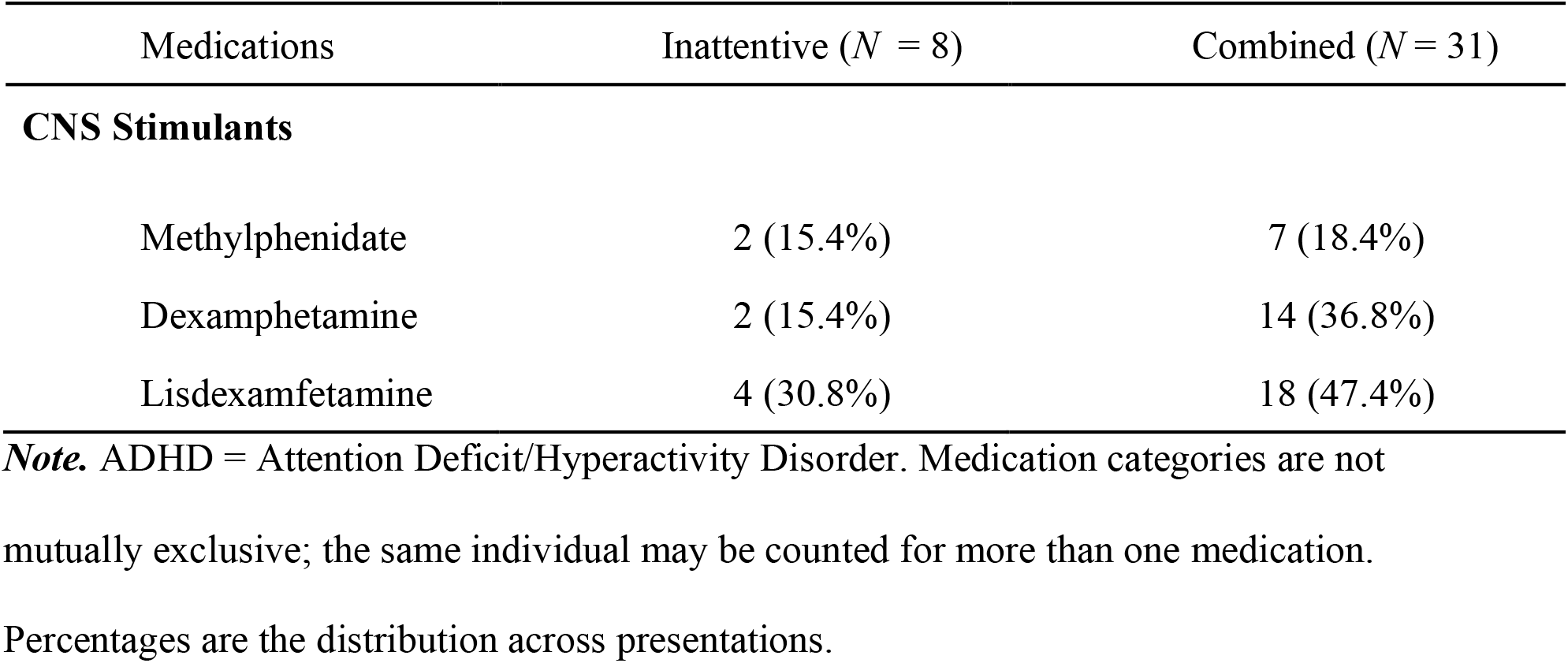
ADHD Medication Use by DIVA Interview Diagnosis.

**Supplementary Table 3.**
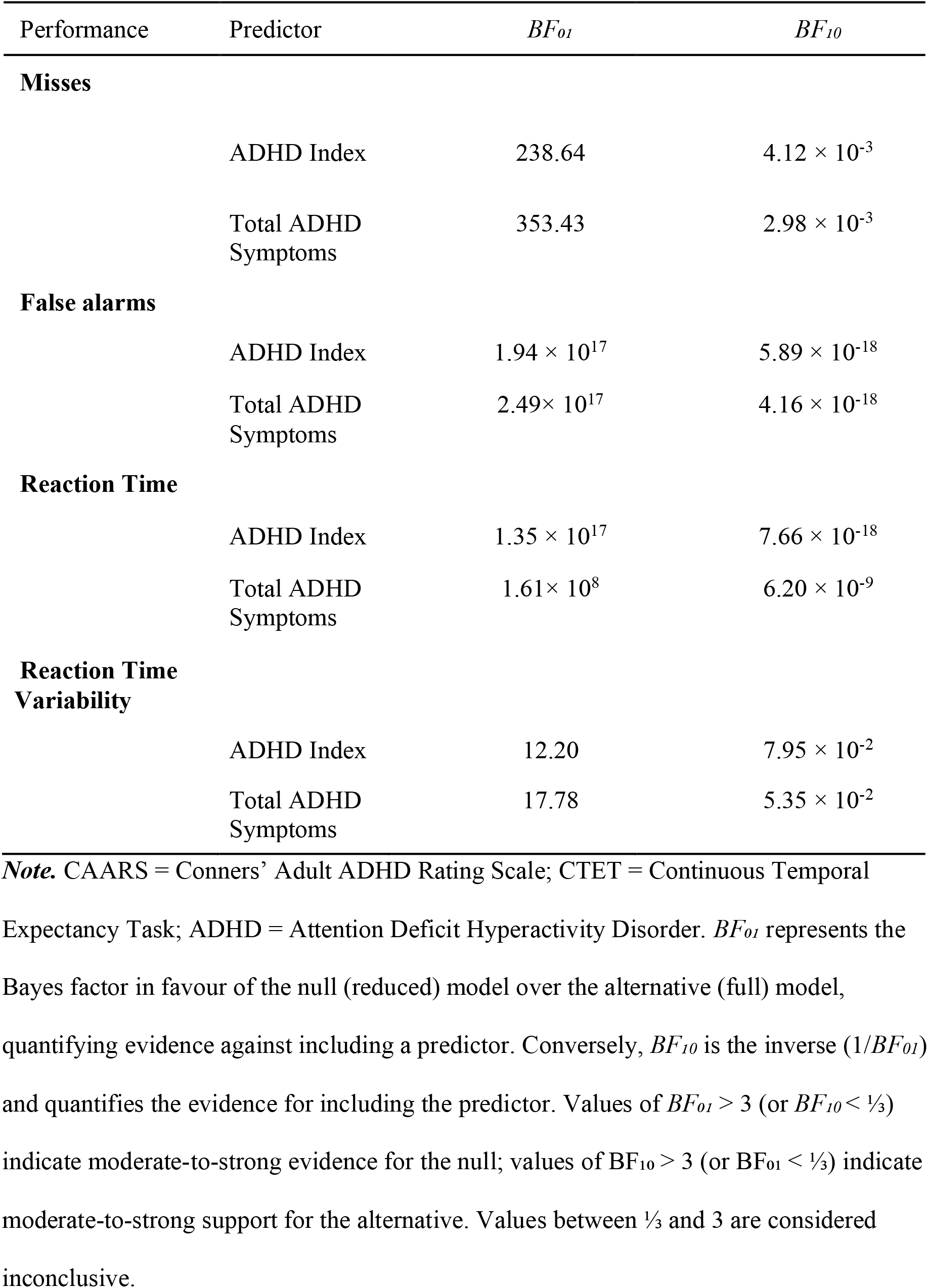
Bayesian model comparisons for the inclusion of CAARS ADHD Index and Total Symptoms across CTET performance measures.

| Performance | Predictor | $BF_{01}$ | $BF_{10}$ |
| --- | --- | --- | --- |
| **Misses** |  |  |  |
| | ADHD Index | 238.64 | $4.12 \times 10^{-3}$ |
| | Total ADHD Symptoms | 353.43 | $2.98 \times 10^{-3}$ |
| **False alarms** |  |  |  |
| | ADHD Index | $1.94 \times 10^{17}$ | $5.89 \times 10^{-18}$ |
| | Total ADHD Symptoms | $2.49 \times 10^{17}$ | $4.16 \times 10^{-18}$ |
| **Reaction Time** |  |  |  |
| | ADHD Index | $1.35 \times 10^{17}$ | $7.66 \times 10^{-18}$ |
| | Total ADHD Symptoms | $1.61 \times 10^8$ | $6.20 \times 10^{-9}$ |
| **Reaction Time Variability** |  |  |  |
| | ADHD Index | 12.20 | $7.95 \times 10^{-2}$ |
| | Total ADHD Symptoms | 17.78 | $5.35 \times 10^{-2}$ |

**Note.** CAARS = Conners' Adult ADHD Rating Scale; CTET = Continuous Temporal Expectancy Task; ADHD = Attention Deficit Hyperactivity Disorder. $BF_{01}$ represents the Bayes factor in favour of the null (reduced) model over the alternative (full) model, quantifying evidence against including a predictor. Conversely, $BF_{10}$ is the inverse ( $1/BF_{01}$ ) and quantifies the evidence for including the predictor. Values of $BF_{01} > 3$ (or $BF_{10} < 1/3$ ) indicate moderate-to-strong evidence for the null; values of $BF_{10} > 3$ (or $BF_{01} < 1/3$ ) indicate moderate-to-strong support for the alternative. Values between $1/3$ and 3 are considered inconclusive.

## Supplementary Figures

**Supplementary Figure 1.**
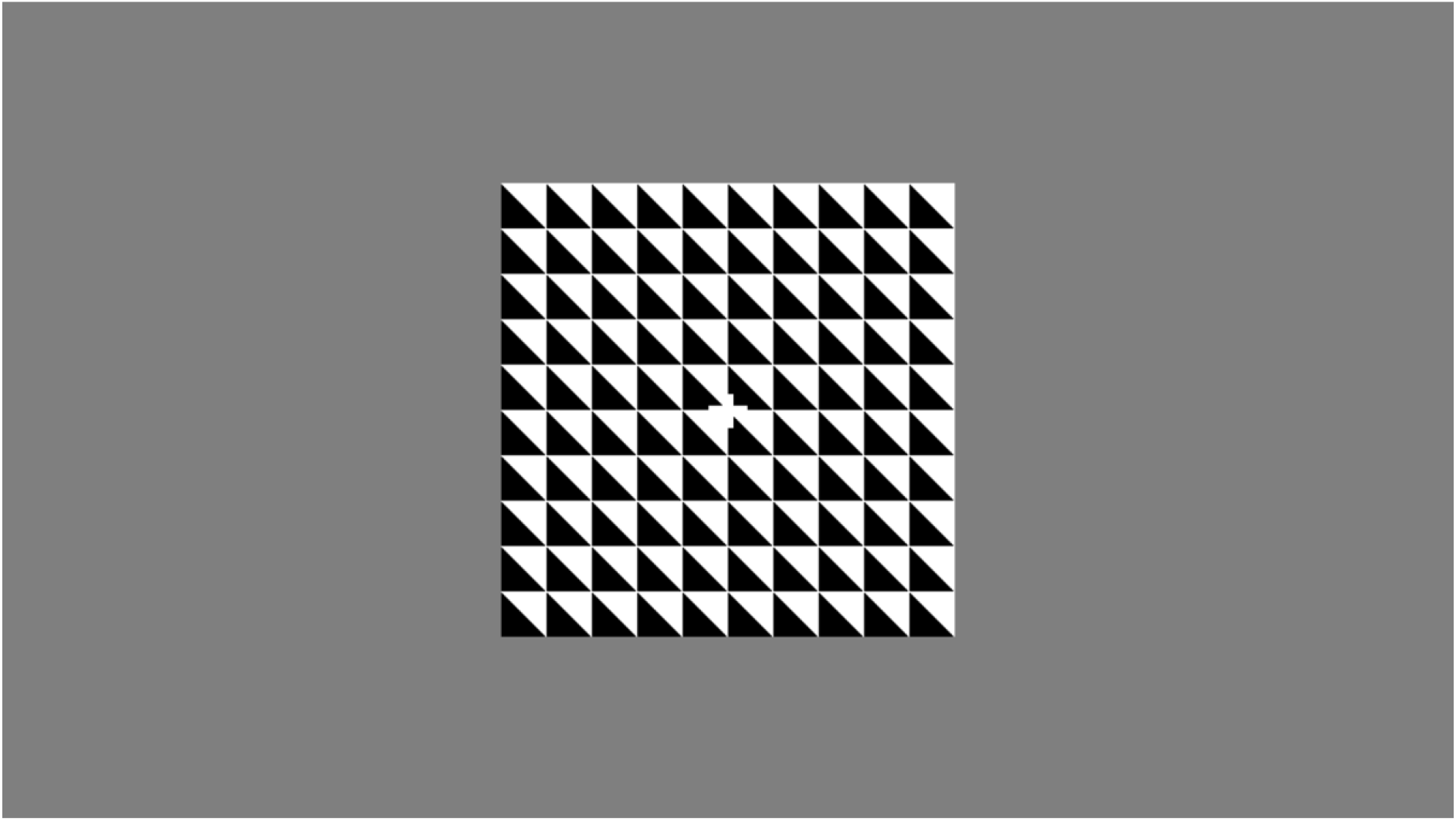
***Note.*** Animated depiction of the Continuous Temporal Expectancy Task (CTET). Standard, non-target stimuli were present for 800 ms and target stimuli for 1120 ms. Stimulus orientation rotated 90° between presentations. In the task, 7–15 non-target stimuli were pseudo-randomly presented between targets (5.6–12 s inter-target interval). The 25 Hz flicker used to elicit steady-state visually evoked potentials (SSVEPs) is not depicted.

**Supplementary Figure 2.**
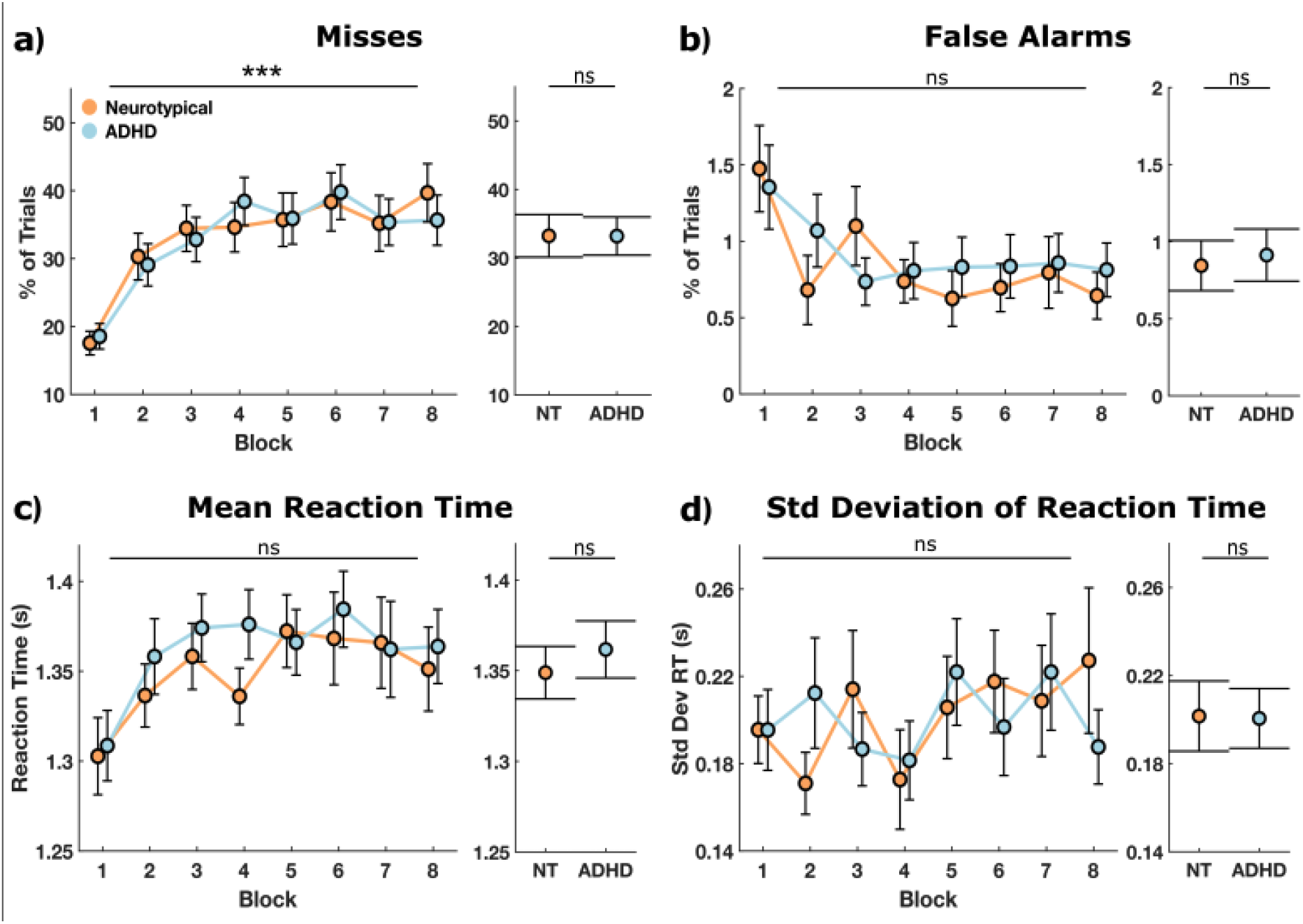
CTET Performance Following Exclusion of Low-Accuracy Participants ***Note***. **a.** Continuous Temporal Expectancy Task (CTET) performance results after excluding participants who missed more than 35% of target trials in the first block (final N: ADHD = 41, Neurotypical (NT) = 35). This threshold was based on performance statistics from the original CTET study (O’Connell et al., 2009), which reported a mean correct detection rate of 64%, suggesting that participants below this threshold were outliers in sustained attention. Panels show: **a.** misses, **b**. false alarms, **c**. mean reaction time, and **e.** standard deviation of reaction time. Left panels show group mean performance (NT in orange and ADHD in blue) across eight blocks, with error bars indicating standard error. Right panels show overall group means and standard error. Stars indicate the strength of the block effect (left panels) and group differences (right panels) as determined by Bayesian linear mixed-effect models (ns: BF₁₀ <⅓, evidence for null; ***BF₁₀ > 30, strong evidence). No differences were found compared to the full-task performance analysis: group differences remained non-significant, and only misses had a block effect.

## References

Andrillon, T., Burns, A., Mackay, T., Windt, J., & Tsuchiya, N. (2021). Predicting lapses of attention with sleep-like slow waves. Nature Communications, 12(1), 3657. 10.1038/s41467-021-23890-7

Andrillon, T., Windt, J., Silk, T., Drummond, S. P. A., Bellgrove, M. A., & Tsuchiya, N. (2019). Does the Mind Wander When the Brain Takes a Break? Local Sleep in Wakefulness, Attentional Lapses and Mind-Wandering. Frontiers in Neuroscience, 13, 949. 10.3389/fnins.2019.00949

Beebe, D. W. (2011). Cognitive, Behavioral, and Functional Consequences of Inadequate Sleep in Children and Adolescents. Pediatric Clinics of North America, 58(3), 649–665. 10.1016/j.pcl.2011.03.002

Bellgrove, M. A., Hawi, Z., Kirley, A., Gill, M., & Robertson, I. H. (2005). Dissecting the attention deficit hyperactivity disorder (ADHD) phenotype: Sustained attention, response variability and spatial attentional asymmetries in relation to dopamine transporter (DAT1) genotype. Neuropsychologia, 43(13), 1847–1857. 10.1016/j.neuropsychologia.2005.03.011

Bernardi, G., Siclari, F., Yu, X., Zennig, C., Bellesi, M., Ricciardi, E., Cirelli, C., Ghilardi, M. F., Pietrini, P., & Tononi, G. (2015). Neural and behavioral correlates of extended training during sleep deprivation in humans: Evidence for local, task-specific effects. The Journal of Neuroscience: The Official Journal of The Society for Neuroscience, 35(11), 4487–4500. (25788668). 10.1523/JNEUROSCI.4567-14.2015

Biabani, M., Walsh, K., Zhou, S.-H., Wagner, J., Johnstone, A., Paterson, J., Johnson, B. P., Matthews, N., Loughnane, G. M., O’Connell, R. G., & Bellgrove, M. A. (2025). Neurophysiology of Perceptual Decision-Making and Its Alterations in Attention-Deficit Hyperactivity Disorder. The Journal of Neuroscience, 45(14), e0469242025. 10.1523/jneurosci.0469-24.2025

Bioulac, S., Micoulaud-Franchi, J.-A., & Philip, P. (2015). Excessive Daytime Sleepiness in Patients With ADHD—Diagnostic and Management Strategies. Current Psychiatry Reports, 17(8), 69. 10.1007/s11920-015-0608-7

Bjorvatn, B., Brevik, E. J., Lundervold, A. J., Halmøy, A., Posserud, M.-B., Instanes, J. T., & Haavik, J. (2017). Adults with Attention Deficit Hyperactivity Disorder Report High Symptom Levels of Troubled Sleep, Restless Legs, and Cataplexy. Frontiers in Psychology, 8. 10.3389/fpsyg.2017.01621

Bozhilova, N., Cooper, R., Kuntsi, J., Asherson, P., & Michelini, G. (2020). Electrophysiological correlates of spontaneous mind wandering in attention-deficit/hyperactivity disorder. Behavioural Brain Research, 391, 112632. 10.1016/j.bbr.2020.112632

Choong, S. Y., Byrne, J. E. M., Drummond, S. P. A., Rispoli-Yovanovic, M., Jones, A., Lum, J. A. G., & Staiger, P. K. (2025). A meta-analytic investigation of the effect of sleep deprivation on inhibitory control. Sleep Medicine Reviews, 80, 102042. 10.1016/j.smrv.2024.102042

Connell, R. G., Dockree, P. M., Robertson, I. H., Bellgrove, M. A., Foxe, J. J., & Kelly, S. P. (2009). Uncovering the neural signature of lapsing attention: Electrophysiological signals predict errors up to 20 s before they occur. The Journal of Neuroscience, 29(26), 8604–8611. (19571151). 10.1523/JNEUROSCI.5967-08.2009

Conners, C. K., Erhardt, D., Epstein, J. N., Parker, J. D. A., Sitarenios, G., & Sparrow, E. (1999). Self-ratings of ADHD symptoms in adults I: Factor structure and normative data. Journal of Attention Disorders, 3(3), 141–151. 10.1177/108705479900300303

Cortese, S., Brown, T. E., Corkum, P., Gruber, R., O’Brien, L. M., Stein, M., Weiss, M., & Owens, J. (2013). Assessment and Management of Sleep Problems in Youths With Attention-Deficit/Hyperactivity Disorder. Journal of the American Academy of Child & Adolescent Psychiatry, 52(8), 784–796. 10.1016/j.jaac.2013.06.001

Cortese, S., Kelly, C., Chabernaud, C., Proal, E., Di Martino, A., Milham, M. P., & Castellanos, F. X. (2012). Toward Systems Neuroscience of ADHD: A Meta-Analysis of 55 fMRI Studies. American Journal of Psychiatry, 169(10), 1038–1055. 10.1176/appi.ajp.2012.11101521

Coull, J., & Nobre, A. (2008). Dissociating explicit timing from temporal expectation with fMRI. Current Opinion in Neurobiology, 18(2), 137–144. 10.1016/j.conb.2008.07.011

Crowley, K. (2002). The effects of normal aging on sleep spindle and K-complex production. Clinical Neurophysiology, 113(10), 1615–1622. 10.1016/s1388-2457(02)00237-7

Czisch, M., Wehrle, R., Harsay, H. A., Wetter, T. C., Holsboer, F., Sämann, P. G., & Drummond, S. P. A. (2012). On the need of objective vigilance monitoring: Effects of sleep loss on target detection and task-negative activity using combined EEG/fMRI. Frontiers in Neurology, 3(67). 10.3389/fneur.2012.00067

Dijk, D.-J. (2009). Regulation and Functional Correlates of Slow Wave Sleep. Journal of Clinical Sleep Medicine, *5*(2 suppl). 10.5664/jcsm.5.2S.S6

Drummond, S. P. A., Anderson, D. E., Straus, L. D., Vogel, E. K., & Perez, V. B. (2012). The Effects of Two Types of Sleep Deprivation on Visual Working Memory Capacity and Filtering Efficiency. PLoS ONE, 7(4), e35653. 10.1371/journal.pone.0035653

Drummond, S. P. A., Paulus, M. P., & Tapert, S. F. (2006). Effects of two nights sleep deprivation and two nights recovery sleep on response inhibition. Journal of Sleep Research, 15(3), 261–265. 10.1111/j.1365-2869.2006.00535.x

Esterman, M., Poole, V., Liu, G., & DeGutis, J. (2016). Modulating Reward Induces Differential Neurocognitive Approaches to Sustained Attention. Cerebral Cortex, cercor;bhw214v1. 10.1093/cercor/bhw214

Faraone, S. V., Bellgrove, M. A., Brikell, I., Cortese, S., Hartman, C. A., Hollis, C., Newcorn, J. H., Philipsen, A., Polanczyk, G. V., Rubia, K., Sibley, M. H., & Buitelaar, J. K. (2024). Attention-deficit/hyperactivity disorder. Nature Reviews Disease Primers, 10(1), 11. 10.1038/s41572-024-00495-0

Fortenbaugh, F. C., DeGutis, J., & Esterman, M. (2017). Recent theoretical, neural, and clinical advances in sustained attention research. Annals of the New York Academy of Sciences, 1396(1), 70–91. 10.1111/nyas.13318

Gobin, C. M., Banks, J. B., Fins, A. I., & Tartar, J. L. (2015). Poor sleep quality is associated with a negative cognitive bias and decreased sustained attention. Journal of Sleep Research, 24(5), 535–542. 10.1111/jsr.12302

Gruber, R., Cassoff, J., Frenette, S., Wiebe, S., & Carrier, J. (2012). Impact of Sleep Extension and Restriction on Children’s Emotional Lability and Impulsivity. Pediatrics, 130(5), e1155–e1161. 10.1542/peds.2012-0564

Helfer, B., Bozhilova, N., Cooper, R. E., Douzenis, J. I., Maltezos, S., & Asherson, P. (2020). The key role of daytime sleepiness in cognitive functioning of adults with attention deficit hyperactivity disorder. European Psychiatry, 63(1), e31. 10.1192/j.eurpsy.2020.28

Hung, C. S., Sarasso, S., Ferrarelli, F., Riedner, B., Ghilardi, M. F., Cirelli, C., & Tononi, G. (2013). Local experience-dependent changes in the wake EEG after prolonged wakefulness. Sleep, 36(1), 59–72. (23288972). 10.5665/sleep.2302

Hvolby, A. (2015). Associations of sleep disturbance with ADHD: Implications for treatment. ADHD Attention Deficit and Hyperactivity Disorders, 7(1), 1–18. 10.1007/s12402-014-0151-0

Kaiser, A., Aggensteiner, P.-M., Baumeister, S., Holz, N. E., Banaschewski, T., & Brandeis, D. (2020). Earlier versus later cognitive event-related potentials (ERPs) in attention-deficit/hyperactivity disorder (ADHD): A meta-analysis. Neuroscience & Biobehavioral Reviews, 112, 117–134. 10.1016/j.neubiorev.2020.01.019

Kliger Amrani, A., & Zion Golumbic, E. (2020). Spontaneous and stimulus-driven rhythmic behaviors in ADHD adults and controls. Neuropsychologia, 146, 107544. 10.1016/j.neuropsychologia.2020.107544

Krause, A. J., Simon, E. B., Mander, B. A., Greer, S. M., Saletin, J. M., Goldstein-Piekarski, A. N., & Walker, M. P. (2017). The sleep-deprived human brain. Nature Reviews Neuroscience, 18(7), 404–418. 10.1038/nrn.2017.55

Lazar, A. S., Lazar, Z. I., & Dijk, D.-J. (2015). Circadian regulation of slow waves in human sleep: Topographical aspects. NeuroImage, 116, 123–134. 10.1016/j.neuroimage.2015.05.012

Liechti, M. D., Valko, L., Müller, U. C., Döhnert, M., Drechsler, R., Steinhausen, H.-C., & Brandeis, D. (2013). Diagnostic Value of Resting Electroencephalogram in Attention-Deficit/Hyperactivity Disorder Across the Lifespan. Brain Topography, 26(1), 135–151. 10.1007/s10548-012-0258-6

Lo, J. C., Groeger, J. A., Santhi, N., Arbon, E. L., Lazar, A. S., Hasan, S., Von Schantz, M., Archer, S. N., & Dijk, D.-J. (2012). Effects of Partial and Acute Total Sleep Deprivation on Performance across Cognitive Domains, Individuals and Circadian Phase. PLoS ONE, 7(9), e45987. 10.1371/journal.pone.0045987

Loo, S. K., Hale, T. S., Macion, J., Hanada, G., McGough, J. J., McCracken, J. T., & Smalley, S. L. (2009). Cortical activity patterns in ADHD during arousal, activation and sustained attention. Neuropsychologia, 47(10), 2114–2119. 10.1016/j.neuropsychologia.2009.04.013

Ma, N., Dinges, D. F., Basner, M., & Rao, H. (2015). How Acute Total Sleep Loss Affects the Attending Brain: A Meta-Analysis of Neuroimaging Studies. Sleep, 38(2), 233–240. 10.5665/sleep.4404

Machida, K., Barry, E., Mulligan, A., Gill, M., Robertson, I. H., Lewis, F. C., Green, B., Kelly, S. P., Bellgrove, M. A., & Johnson, K. A. (2022). Which Measures From a Sustained Attention Task Best Predict ADHD Group Membership? Journal of Attention Disorders, 26(11), 1471–1482. 10.1177/10870547221081266

Madiouni, C., Lopez, R., Gély-Nargeot, M.-C., Lebrun, C., & Bayard, S. (2020). Mind-wandering and sleepiness in adults with attention-deficit/hyperactivity disorder. Psychiatry Research, 287, 112901. 10.1016/j.psychres.2020.112901

Maris, E., & Oostenveld, R. (2007). Nonparametric statistical testing of EEG- and MEG-data. Journal of Neuroscience Methods, 164(1), 177–190. 10.1016/j.jneumeth.2007.03.024

Markovska-Simoska, S., & Pop-Jordanova, N. (2017). Quantitative EEG in Children and Adults With Attention Deficit Hyperactivity Disorder: Comparison of Absolute and Relative Power Spectra and Theta/Beta Ratio. Clinical EEG and Neuroscience, 48(1), 20–32. 10.1177/1550059416643824

May, T., Birch, E., Chaves, K., Cranswick, N., Culnane, E., Delaney, J., Derrick, M., Eapen, V., Edlington, C., Efron, D., Ewais, T., Garner, I., Gathercole, M., Jagadheesan, K., Jobson, L., Kramer, J., Mack, M., Misso, M., Murrup-Stewart, C., … Bellgrove, M. (2023). The Australian evidence-based clinical practice guideline for attention deficit hyperactivity disorder. Australian & New Zealand Journal of Psychiatry, 57(8), 1101–1116. 10.1177/00048674231166329

Morgan, S. T., Hansen, J. C., & Hillyard, S. A. (1996). Selective attention to stimulus location modulates the steady-state visual evoked potential. Proceedings of the National Academy of Sciences, 93(10), 4770–4774. 10.1073/pnas.93.10.4770

Müller, M. M., Picton, T. W., Valdes-Sosa, P., Riera, J., Teder-Sälejärvi, W. A., & Hillyard, S. A. (1998). Effects of spatial selective attention on the steady-state visual evoked potential in the 20–28 Hz range. Cognitive Brain Research, 6(4), 249–261. 10.1016/S0926-6410(97)00036-0

O’Connell, R. G., Dockree, P. M., Robertson, I. H., Bellgrove, M. A., Foxe, J. J., & Kelly, S. P. (2009). Uncovering the Neural Signature of Lapsing Attention: Electrophysiological Signals Predict Errors up to 20 s before They Occur. Journal of Neuroscience, 29(26), 8604–8611. 10.1523/JNEUROSCI.5967-08.2009

Oostenveld, R., Fries, P., Maris, E., & Schoffelen, J.-M. (2011). FieldTrip: Open Source Software for Advanced Analysis of MEG, EEG, and Invasive Electrophysiological Data. Computational Intelligence and Neuroscience, 2011, 1–9. 10.1155/2011/156869

Pinggal, E., Dockree, P. M., O’Connell, R. G., Bellgrove, M. A., & Andrillon, T. (2022). Pharmacological manipulations of physiological arousal and sleep-like slow waves modulate sustained attention. Journal of Neuroscience, 8113–8124. 10.1523/JNEUROSCI.0836-22.2022

Pinggal, E., Jackson, J., Kusztor, A., Chapman, D., Windt, J., Drummond, S. P. A., Silk, T. J., Bellgrove, M. A., & Andrillon, T. (2026). Sleep-like slow waves during wakefulness mediate attention and vigilance difficulties in adult attention-deficit/hyperactivity disorder. The Journal of Neuroscience, e1694252025. 10.1523/JNEUROSCI.1694-25.2025

Polich, J. (2007). Updating P300: An integrative theory of P3a and P3b. Clinical Neurophysiology, 118(10), 2128–2148. 10.1016/j.clinph.2007.04.019

Quercia, A., Zappasodi, F., Committeri, G., & Ferrara, M. (2018). Local use-dependent sleep in wakefulness links performance errors to learning. Frontiers in Human Neuroscience, 12, 122. PubMed (29666574). 10.3389/fnhum.2018.00122

Raffaelli, Q., Rai, S., Galbraith, A., Krupa, A., Buerkner, J., Andrews-Hanna, J. R., Callahan, B. L., & Kam, J. W. Y. (2025). Hyperactive ADHD symptoms are associated with increased variability in thought content in less constrained contexts. Scientific Reports, 15(1), 9792. 10.1038/s41598-025-93053-x

Riedner, B. A., Vyazovskiy, V. V., Huber, R., Massimini, M., Esser, S., Murphy, M., & Tononi, G. (2007). Sleep homeostasis and cortical synchronization: III. A high-density EEG study of sleep slow waves in humans. Sleep, 30(12), 1643–1657. 10.1093/sleep/30.12.1643

Robertson, I. H., Manly, T., Andrade, J., Baddeley, B. T., & Yiend, J. (1997). Sustained Attention to Response Task [Dataset]. 10.1037/t28308-000

Saletin, J. M., Jackvony, S., Rodriguez, K. A., & Dickstein, D. P. (2019). A coordinate-based meta-analysis comparing brain activation between attention deficit hyperactivity disorder and total sleep deprivation. Sleep, 42(3), zsy251. 10.1093/sleep/zsy251

Smallwood, J., Davies, J. B., Heim, D., Finnigan, F., Sudberry, M., O’Connor, R., & Obonsawin, M. (2004). Subjective experience and the attentional lapse: Task engagement and disengagement during sustained attention. Consciousness and Cognition, 13(4), 657–690. 10.1016/j.concog.2004.06.003

Steriade, M. (2005). Sleep, epilepsy and thalamic reticular inhibitory neurons. Trends Neurosci, 28(6), 317–324. (15927688). 10.1016/j.tins.2005.03.007

Szuromi, B., Czobor, P., Komlósi, S., & Bitter, I. (2011). P300 deficits in adults with attention deficit hyperactivity disorder: A meta-analysis. Psychological Medicine, 41(7), 1529–1538. 10.1017/S0033291710001996

Ustun, B., Adler, L. A., Rudin, C., Faraone, S. V., Spencer, T. J., Berglund, P., Gruber, M. J., & Kessler, R. C. (2017). The World Health Organization Adult Attention-Deficit/Hyperactivity Disorder Self-Report Screening Scale for *DSM-5*. JAMA Psychiatry, 74(5), 520. 10.1001/jamapsychiatry.2017.0298

Van Den Driessche, C., Bastian, M., Peyre, H., Stordeur, C., Acquaviva, É., Bahadori, S., Delorme, R., & Sackur, J. (2017). Attentional Lapses in Attention-Deficit/Hyperactivity Disorder: Blank Rather Than Wandering Thoughts. Psychological Science, 28(10), 1375–1386. 10.1177/0956797617708234

Vlahou, E. L., Thurm, F., Kolassa, I.-T., & Schlee, W. (2014). Resting-state slow wave power, healthy aging and cognitive performance. Scientific Reports, 4(1). 10.1038/srep05101

Vyazovskiy, V. V., Olcese, U., Hanlon, E. C., Nir, Y., Cirelli, C., & Tononi, G. (2011). Local sleep in awake rats. Nature, 472(7344), 443–447. (21525926). 10.1038/nature10009

