## Supplementary Tables and Figures for "Diminished Stimulus-Evoked Activity During Sustained Attention Without Behavioural Cost in ADHD"

**Supplementary Table 1.**

*Demographics and Questionnaire Data*

|  | ADHD (*N* = 51) | Neurotypical (*N* = 48) | *p*-Value |
| --- | --- | --- | --- |
| **Wender Utah Rating Scale** (mean (SD)) | 58 (15.9) |  |  |
| **Inventory of Depression and Anxiety Symptoms (IDAS-II)** (mean (SD)) |  |  |  |
| General Depression | 60.7 (12.9) | 36.9 (10.9) | < .001 |
| Dysphoria | 30.3 (7.4) | 18.0 (6.6) | < .001 |
| Well Being | 20.8 (6.2) | 24.9 (6.5) | .002 |
| Panic | 14.3 (5.1) | 9.9 (3.5) | < .001 |
| **Alcohol Use Disorders Identification Test** (mean (SD)) | 5.2 (4.7) | 2.7 (2.4) | .001 |
| **Drug Use Disorders Identification Test** (mean (SD)) | 3.3 (5.8) | 0.4 (1.0) | .001 |
| **National Adult Reading Test (NART)** | 114.2 (3.6) | 110.9 (6.9) | .004 |

***Note.*** ADHD = Attention Deficit/Hyperactivity Disorder. Higher WURS scores reflect greater retrospectively reported childhood ADHD symptoms. Higher scores on the IDAS-II subscales reflect greater symptom severity within each domain. Higher AUDIT and DUDIT scores reflect greater alcohol consumption and drug use, respectively. Higher NART scores indicate higher estimated verbal IQ. Between-group comparisons were conducted using Welch’s *t*-tests, with corresponding *p-*values reported.

**Supplementary Table 2.**

*ADHD Medication Use by DIVA Interview Diagnosis*

| Medications | Inattentive (*N*  = 8) | Combined (*N* = 31) |
| --- | --- | --- |
| **CNS** **Stimulants** |  |  |
| Methylphenidate | 2 (15.4%) | 7 (18.4%) |
| Dexamphetamine | 2 (15.4%) | 14 (36.8%) |
| Lisdexamfetamine | 4 (30.8%) | 18 (47.4%) |

***Note.*** ADHD = Attention Deficit/Hyperactivity Disorder. Medication categories are not mutually exclusive; the same individual may be counted for more than one medication. Percentages are the distribution across presentations.

**Supplementary Table 3.**

*Bayesian model comparisons for the inclusion of CAARS ADHD Index and Total Symptoms across CTET performance measures*

| Performance | Predictor | *BF₀₁* | *BF₁_0_* |
| --- | --- | --- | --- |
| **Misses** |  |  |  |
|  | ADHD Index | 238.64 | 4.12 × 10^-3^ |
|  | Total ADHD Symptoms | 353.43 | 2.98 × 10^-3^ |
| **False alarms** |  |  |  |
|  | ADHD Index | 1.94 × 10^17^ | 5.89 × 10^-18^ |
|  | Total ADHD Symptoms | 2.49× 10^17^ | 4.16 × 10^-18^ |
| **Reaction Time** |  |  |  |
|  | ADHD Index | 1.35 × 10^17^ | 7.66 × 10^-18^ |
|  | Total ADHD Symptoms | 1.61× 10^8^ | 6.20 × 10^-9^ |
| **Reaction Time Variability** |  |  |  |
|  | ADHD Index | 12.20 | 7.95 × 10^-2^ |
|  | Total ADHD Symptoms | 17.78 | 5.35 × 10^-2^ |

***Note.*** CAARS = Conners’ Adult ADHD Rating Scale; CTET = Continuous Temporal Expectancy Task; ADHD = Attention Deficit Hyperactivity Disorder. *BF₀₁* represents the Bayes factor in favour of the null (reduced) model over the alternative (full) model, quantifying evidence against including a predictor. Conversely, *BF₁_0_* is the inverse (1/*BF_01_*) and quantifies the evidence for including the predictor. Values of *BF₀₁* > 3 (or *BF₁_0_* < ⅓) indicate moderate-to-strong evidence for the null; values of BF₁₀ > 3 (or BF₀₁ < ⅓) indicate moderate-to-strong support for the alternative. Values between ⅓ and 3 are considered inconclusive.

**Supplementary Figures**

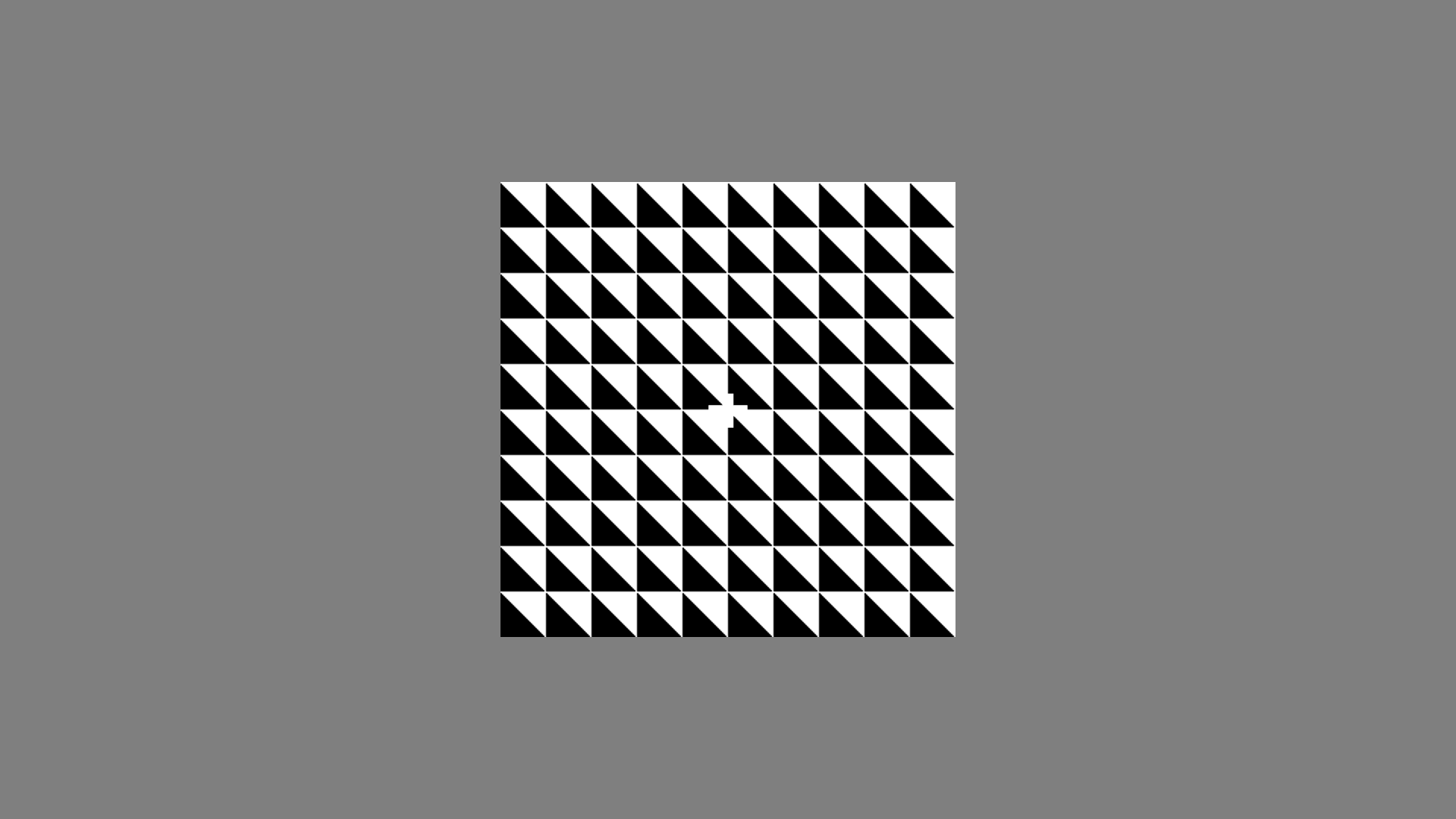

**Supplementary Figure 1.**

***Note.*** Animated depiction of the Continuous Temporal Expectancy Task (CTET). Standard, non-target stimuli were present for 800 ms and target stimuli for 1120 ms. Stimulus orientation rotated 90° between presentations. In the task, 7–15 non-target stimuli were pseudo-randomly presented between targets (5.6–12 s inter-target interval). The 25 Hz flicker used to elicit steady-state visually evoked potentials (SSVEPs) is not depicted.

**Supplementary Figure 2.**

*CTET Performance Following Exclusion of Low-Accuracy Participants*
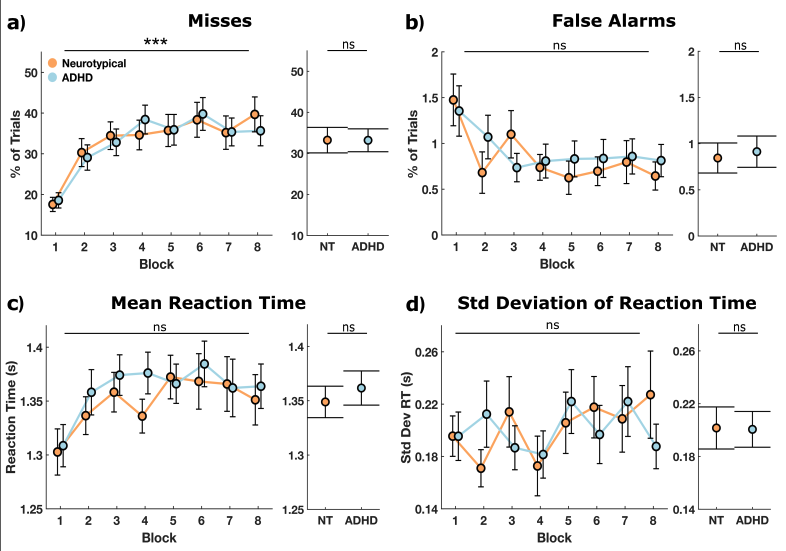

***Note***. **a.** Continuous Temporal Expectancy Task (CTET) performance results after excluding participants who missed more than 35% of target trials in the first block (final N: ADHD = 41, Neurotypical (NT) = 35). This threshold was based on performance statistics from the original CTET study (O’Connell et al., 2009), which reported a mean correct detection rate of 64%, suggesting that participants below this threshold were outliers in sustained attention. Panels show: **a.** misses, **b**. false alarms, **c**. mean reaction time, and **e.** standard deviation of reaction time. Left panels show group mean performance (NT in orange and ADHD in blue) across eight blocks, with error bars indicating standard error. Right panels show overall group means and standard error. Stars indicate the strength of the block effect (left panels) and group differences (right panels) as determined by Bayesian linear mixed-effect models (ns: BF₁₀ <⅓, evidence for null; ***BF₁₀ > 30, strong evidence). No differences were found compared to the full-task performance analysis: group differences remained non-significant, and only misses had a block effect.
